# Hierarchical Assembly of Native Cytoplasmic Lattices Revealed by Cryo-EM

**DOI:** 10.64898/2026.03.30.715201

**Authors:** Yin-Li Zhang, Yuqi Liu, Qi Liu, Qi Zhang, Jiamin Jin, Siya Liu, Shuo Yang, Yinan Liu, Ye-Jun Peng, Mengru Lai, Yuting Sun, Ying-Yi Zhang, Yuqing Mei, Xiaomei Tong, Songying Zhang, Miao Gui

## Abstract

Oocytes depend on maternal factors for maturation and early embryogenesis. Cytoplasmic lattices (CPLs) are distinctive fibrillar structures in mammalian oocyte cytoplasm implicated in maternal factor storage and developmental competence, yet their molecular composition and architecture remain elusive. Here, we use cryo-electron microscopy to resolve the high-resolution structure of native mouse oocyte CPLs. We identify 14 intrinsic components of multiple copies that assemble into filaments in which PADI6-mediated self-assembly and two subtypes of subcortical maternal complexes (SCMCs) form the core scaffold. Unexpectedly, we discover that α/β-tubulin dimers and two autoinhibited ubiquitination modules (UHRF1-UBE2D3, SKP1-FBXW18) are internal components of CPLs. We also uncover multivalent interactions that organize adjacent CPL filaments into higher-order helical bundles. By integrating CPL structure with biochemical and proteomic data, we propose a hierarchical assembly mechanism of CPLs, from subcomplexes to filaments to supramolecular lattices. Our study offers mechanistic insights into how maternal proteins are spatially organized to ensure successful early embryonic development and provides a structural foundation for understanding female infertility related to CPL-associated gene mutations.

## Introduction

Mammalian oocytes accumulate extensive reservoirs of maternal proteins and RNAs that sustain fertilization and early embryonic development^1–4^. These maternal factors orchestrate key developmental processes, including cytoskeletal reorganization, epigenetic reprogramming, protein homeostasis and the activation of the embryonic genome^3–5^. Central to this regulatory framework are cytoplasmic lattices (CPLs), specialized filamentous assemblies present in mammalian oocytes and preimplantation embryos that organize and stabilize intracellular constituents^1^. Current models propose that CPLs function as a regulated storage platform, safeguarding maternal factors from degradation and enabling their timely deployment, thereby supporting the successful initiation of mammalian development^6^.

Early studies utilizing transmission electron microscopy (TEM), cryo-electron tomography (cryo-ET) and fluorescence light microscopy have characterized the general morphology of CPLs^6,7^, yet their resolution limits have precluded a detailed understanding of the precise molecular organization of these polymers. Genetic approaches have identified several maternal-effect genes in humans – including *PADI6*, *NLRP5*, *TLE6, OOEP,* and *KHDC3L* – as essential for CPL formation, as their mutation disrupts CPL integrity and leads to early embryonic arrest and human infertility^8–12^. While structures of recombinant subcomplexes, such as the subcortical maternal complex (SCMC) and the PADI6 dimer, have provided insights into local architecture^13–16^, the mechanism governing their assembly into a functional lattice remains unknown. Furthermore, ribosomes, RNAs, and intermediate filaments have been implicated in CPLs^1,17^. It is unclear whether they are intrinsic structural components or merely associated, and what additional factors may contribute to CPL formation and function.

Here, we used cryo-EM to determine the structure of mouse oocyte CPLs, identifying fourteen protein components organized into periodic filaments that further assemble into helical bundles. These findings demonstrate that CPLs function as a multifunctional hub for the storage and coordinated regulation of cytoskeletal elements, the protein degradation machinery and epigenetic regulators. Complemented by proteomics and *in vitro* recombinant studies, we propose a model for the hierarchical assembly pathway of CPL filaments.

## Results

### Structure determination of cytoplasmic lattices from mouse oocytes

To determine the high-resolution structure of intact CPLs, we collected mouse oocytes at the germinal vesicle (GV) stage. Because CPLs readily disassemble upon cell disruption, we developed a cell-lysis-on-grid (CLoG) approach, in which 400-500 oocytes were incubated on glow-discharged cryo-EM grids with lysis buffer containing 0.05% lauryl maltose neopentyl glycol (LMNG) (Figure 1A). An integrated particle-picking strategy was employed to select CPL filaments (Figure S1), yielding a consensus reconstruction of the repeating unit with overall C2 symmetry and local asymmetry features. Local three-dimensional (3D) classifications and refinements improved the maps to a resolution of up to 3.4 Å, permitting unambiguous assignment of fourteen proteins and building of their atomic models (Figures 1B-1D, S1-S2, Tables S1-S3, Supplementary Figures S1-S12). The CPL filament features periodic bulbous swellings separated by narrower segments, with an axial repeat of approximately 37 nm. The filament diameter varies from ∼21 nm at the constricted regions to ∼38 nm at the expanded region. These structural features are consistent with the *in situ* cryo-ET structure of CPLs within mouse oocytes^6^, indicating that the native architecture is preserved following the CLoG procedure.

**Figure 1.**
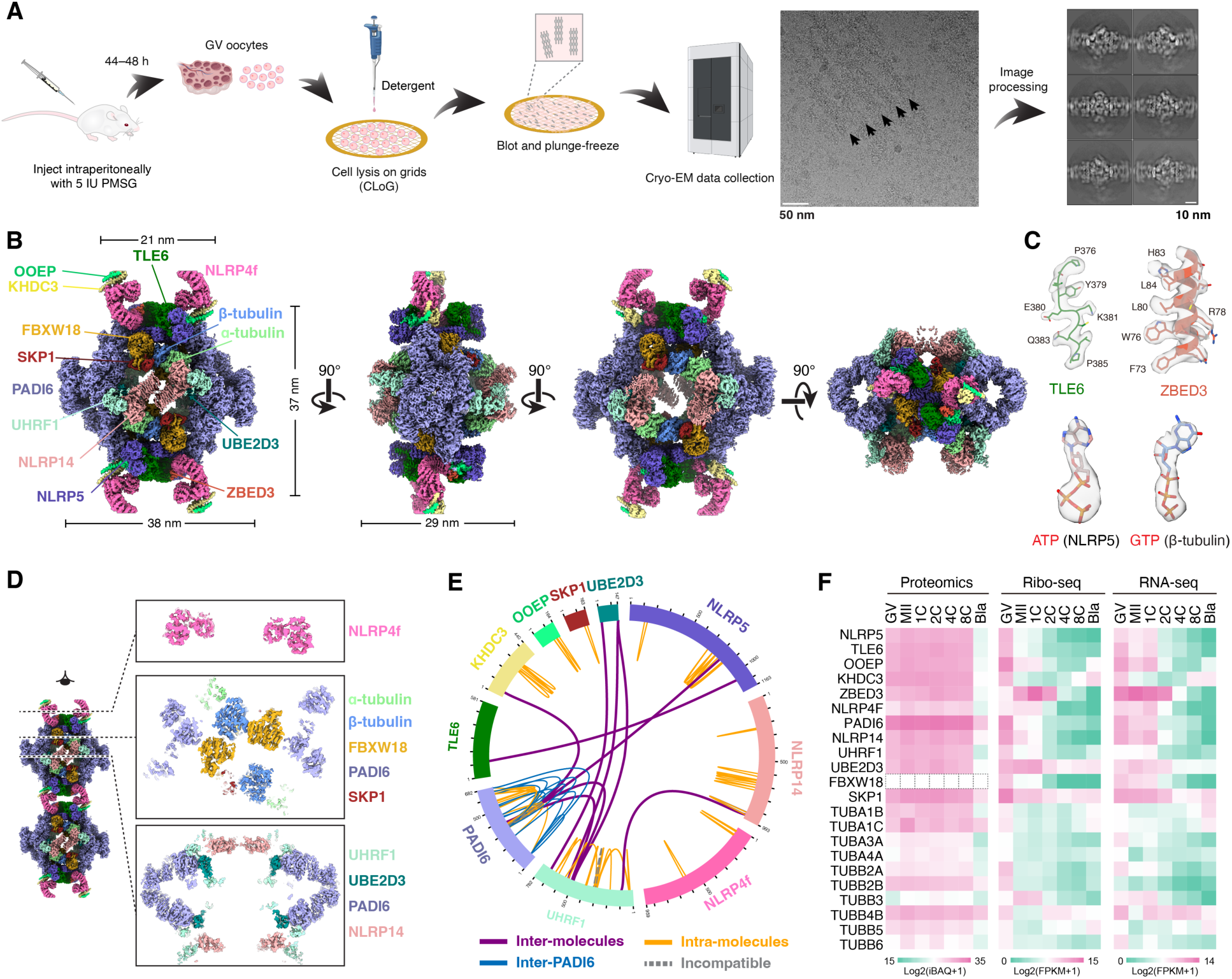
Cryo-EM structure of mouse CPLs. (A) Workflow diagram of the cell-lysis-on-grid (CLoG) method for preparing intact mouse CPLs for cryo-EM. Representative cryo-EM micrographs and 2D class averages are shown. (B) Cryo-EM map of one repeating unit of the CPLs with C2 symmetry. Each protein component is shown in a distinct color. (C) Representative density examples of the cryo-EM reconstruction. (D) Three cross sections of the mouse CPL density map. (E) Diagram of XL-MS analysis of mouse CPLs, highlighting cross-links among PADI6, NLRP5, KHDC3, UHRF1, UBE2D3 and NLRP14. (F) Heatmaps showing the protein, translation and mRNA dynamics of components in CPLs in oocytes and early embryos.

Each bulbous repeat of the CPL filament contains at least ten PADI6 dimers, four SCMCs, four UHRF1-UBE2D3 heterodimers, five SKP1-FBXW18 heterodimers, four NLRP14 molecules and two NLRP4f molecules (Figure 1B). PADI6 and the SCMC (comprising NLRP5, TLE6, OOEP, KHDC3 or ZBED3) form the primary scaffold at the periphery of each bulbous unit. In contrast, functional modules UHRF1-UBE2D3, SKP1-FBXW18, and α/β-tubulin localize to the central core region, whereas NLRP4f bridges adjacent bulbous units to support filament continuity. Together, these components assemble into a densely interconnected molecular network that reinforces the structural integrity of the CPLs (Figure S3A).

The structural organization and protein assignments of CPLs were further validated by cross-linking mass spectrometry (XL-MS), which revealed the close spatial proximity among CPL components at side-chain resolution (Figures 1E, S3C, Tables S4-S5). In total, nine inter-protein cross-links were identified, all of which were fully consistent with the structural model. Specifically, PADI6 peptides were cross-linked to NLRP5, KHDC3, and UHRF1, whereas UHRF1 formed additional cross-links with UBE2D3 and NLRP14. A cross-link was also detected between two SCMC components, NLRP5 and TLE6.

Reanalysis of published proteomic, Ribo-seq, and RNA-seq datasets from mouse oocytes and early embryos spanning developmental stages confirmed high expression of CPL components^18,19^ (Figures 1F and S4). Notably, transcript levels and translational activity decline from high levels at the GV stage to low levels at the two-cell stage, whereas protein abundances remain constant until the eight-cell stage. This temporal pattern is consistent with the persistence of CPL structures from fully grown GV oocytes through early embryonic development^20^.

To further validate protein interactions and identify potential accessory components of CPLs, we performed co-immunoprecipitation (co-IP) coupled with mass spectrometry using mouse oocytes microinjected with cRNAs encoding Flag-tagged *Padi6* or *Ooep* (Figure S5A). A total of 165 and 71 proteins were significantly enriched in the PADI6 and OOEP co-IP datasets, respectively (fold change > 1.5, *p*-value < 0.05) (Figures S5B and S5C), with 42 proteins shared between the two groups, including SKP1 and PADI6 itself (Figure S5D). UHRF1 and UBE2D3 were specifically enriched in the PADI6 interactome, whereas all five SCMC components were enriched in the OOEP group (Figures S5B and S5C). These results support the existence of tightly associated subcomplexes – such as UHRF1-UBE2D3 and the SCMC – that likely function as modular building blocks for the assembly of CPL filaments. Furthermore, we observed a significant enrichment of ribosome proteins within the PADI interactome compared to the OOEP group (fold change > 1.3, p < 0.05) (Figures S5E-S5H). This finding points to a moderate association between ribosome proteins and CPLs facilitated by PADI6, although ribosomes do not appear to be integral components of the lattices.

### PADI6 self-assembly establishes the CPL scaffold

PADI6 (Protein-arginine deiminase type-6) is a well-characterized maternal factor whose mutations cause early embryonic arrest and female infertility in both humans and mice^8,21,22^. Within CPLs, five PADI6 dimers assemble into a hollow tubular structure at each lateral edge of the bulbous unit (Figure 2A). In addition, two flexible PADI6 dimers cap the ends of the tubule and participate in inter-filament interactions (see below). The PADI6 tubule adopts an imperfect helical architecture, with a variable helical rise ranging from 2.6 to 4.1 nm and a helical twist of 90° to 128° (Figure 2B). This assembly is primarily mediated by interactions between the N-terminal (PAD-N) and middle (PAD-M) domains of adjacent dimers (Figures 2C and 2D). Notably, XL-MS revealed multiple intra– and inter-molecular cross-links among PADI6 molecules, supporting their oligomeric organization within CPLs (Figures 1E and S3C). Moreover, the PAD domains of each dimer engage in lateral contacts with those of two neighboring dimers, creating a network of multivalent interactions that may further stabilize the PADI6 tubular scaffold (Figures 2A and 2B).

**Figure 2.**
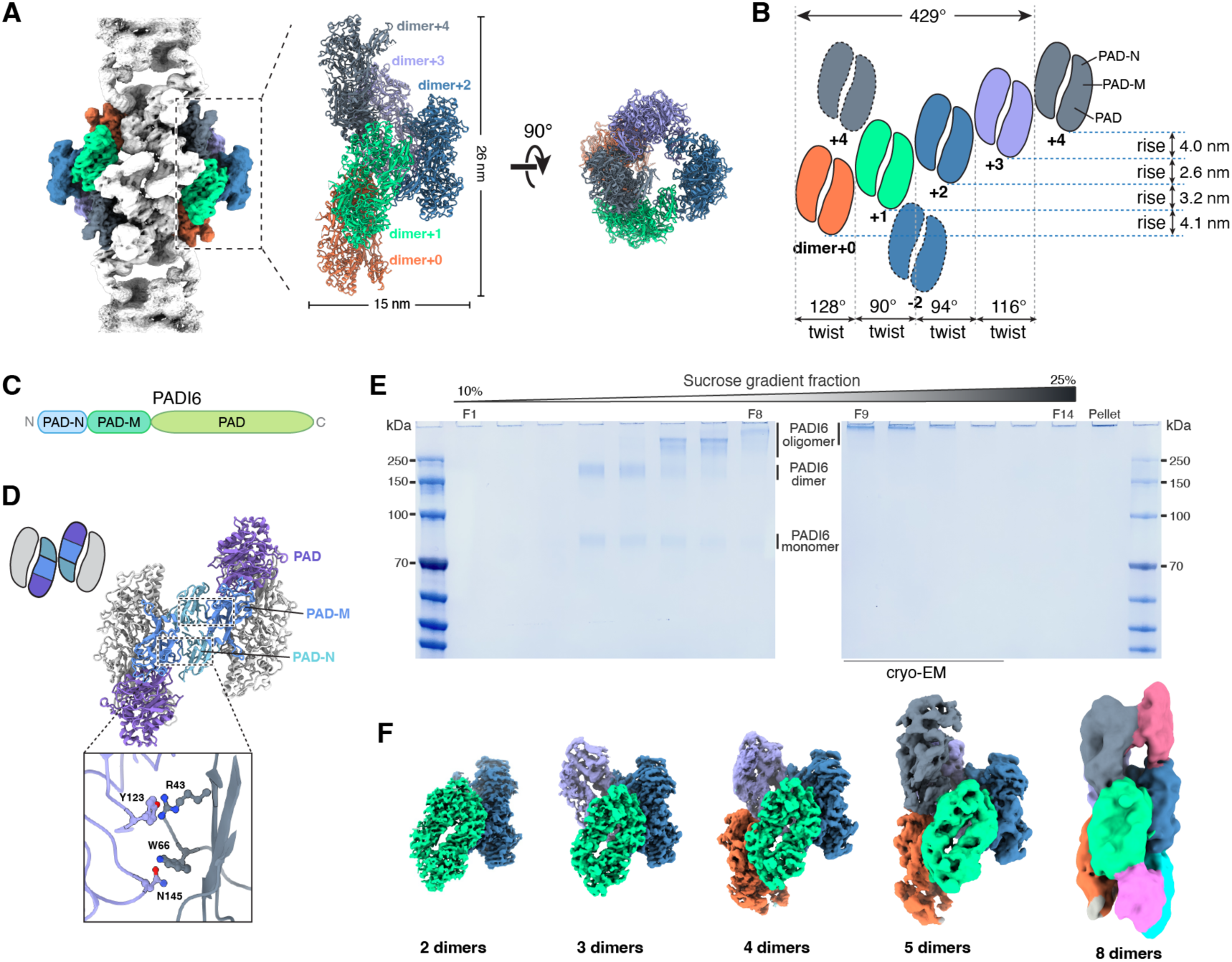
Structural organization and assembly of PADI6. (A) PADI6 dimers within the CPL. The cryo-EM map of the CPL is shown in the left panel, with a low-resolution map outlining the overall assembly. Densities corresponding to the sixth and seventh PADI6 dimer are visible in low contour level map. Model of five PADI6 dimers is shown in the right panel. Each PADI6 dimer is shown with a distinct color. (B) Cartoon schematic illustrating the arrangement of PADI6 dimers within the CPLs. The helical axis is defined by a line passing through the centroid of the PADI6 tubule and oriented along the longitudinal axis of the CPL. (C) Schematic representation of the domain architecture of PADI6. (D) Inter-dimer interactions mediated by the R43-Y123 and W66-N145 residue pairs in the PADI6 tubule. PADI6 domains are shown in distinct colors. (E) SDS-PAGE analysis showing the oligomeric states of PADI6 after GraFix. (F) Cryo-EM maps of different assemblies of recombinant PADI6 dimers. Each PADI6 dimer is shown with a distinct color.

Previous structural studies of human PADI6 have characterized a conserved dimeric architecture^15,16,22^. To investigate whether PADI6 can self-assemble into higher-order structures analogous to those observed in CPLs, we expressed and purified mouse PADI6 in HEK293F cells and analyzed its oligomeric state by sucrose gradient ultracentrifugation in the presence of the cross-linker BS3 (also known as GraFix)^23^ (Figures 2E, S6A and S6B). In addition to the expected monomeric and dimeric species, SDS-PAGE revealed prominent high-molecular-weight bands, indicating the formation of higher-order assemblies (Figure 2E). Cryo-EM analysis revealed a heterogeneous population comprising oligomers of one to eight dimers (Figures 2F and S6). Notably, the inter-dimer assembly packing closely resembles that observed in CPLs, with a more compact arrangement in the central region relative to the terminal dimers (Figure S7A and S7B). The PAD-N domain, which is flexible in isolated PADI6 dimers, becomes ordered upon engaging the PAD-M domain of an adjacent dimer, thereby stabilizing the tubule assembly (Figure S7C).

Specific interactions at the inter-dimer interface – including contacts between R43 and Y123, and between W66 and N145 – mediate tubular assembly, and disruption of these interactions impairs the higher-order oligomerization of PADI6 (Figures 2D, S7E and S7F). In addition, enzymatic assays showed that purified mouse PADI6 lacks detectable catalytic activity (Figure S6C). This observation is consistent with its substantial evolutionary divergence from other PADI family members (Figure S7D) and with previous reports indicating that the catalytic function of PADI6 is dispensable for CPL formation^15,16,24^. Collectively, these findings establish that PADI6 dimers function primarily as a structural scaffold through their intrinsic polymerization capacity, providing the foundational framework for CPL assembly.

### Two different types of SCMCs bridge the CPL assembly

Unexpectedly, our structure revealed two distinct subtypes of SCMCs, one comprising NLRP5, TLE6, OOEP and KHDC3 (referred as SCMC^KHDC3^) and another in which KHDC3 is replaced by ZBED3 (referred as SCMC^ZBED3^) (Figure 3A). Dimerization of SCMC^KHDC3^ and SCMC^ZBED3^ is mediated by the leucine-rich repeats (LRRs) of NLRP5 molecules (Figure 3B), consistent with the architecture observed in recombinant SCMC structures^13^. In SCMC^KHDC3^, the K Homology (KH) domain of OOEP interacts with the N-terminal pyrin domain (PYD) of NLRP4f, while the C-terminal helix of OOEP engages KHDC3 (Figure 3C). A recent cryo-EM study showed that incubation of NLRP5-TLE6-OOEP, the core of both SCMCs, with KHDC3 failed to resolve densities for KHDC3 and the OOEP C-terminal helix, suggesting that the PYD of NLRP4f stabilizes the SCMC^KHDC3^ assembly^13^. Notably, mouse NLRP5 lacks the N-terminal PYD present in its human counterpart. In recombinant human SCMC, the binding site occupied by mouse NLRP4f PYD is instead engaged by the PYD of human NLRP5 and cannot be substituted by human NLRP4 PYD, highlighting species-specific differences in SCMC and CPL assembly^14^. In SCMC^ZBED3^, ZBED3 replaces the NLRP4f PYD to interact with the SCMC core, which further interacts with the C-terminal LRR of NLRP4f (Figure 3D). Thus, two SCMC dimers are bridged by two NLRP4f molecules to form a hollow, centrosymmetric assembly that links adjacent bulbous units and promotes CPL filament formation.

**Figure 3.**
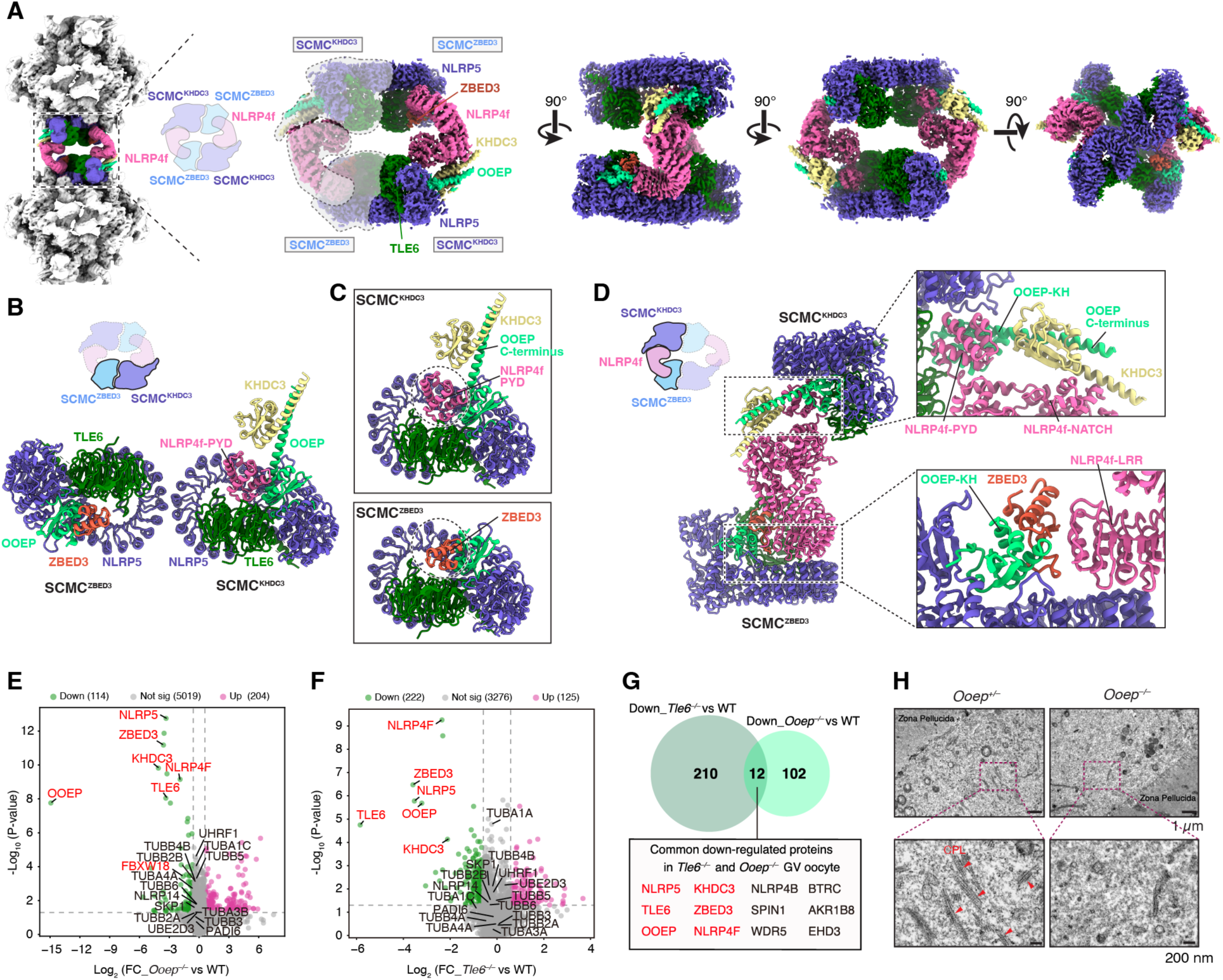
Architecture of the SCMCs within mouse CPLs. (A) Two conformations of SCMC complexes within the CPL, together with NLRP4f, bridge adjacent bulbous units. Each protein is shown in a distinct color. (B) Dimerization of SCMC^KHDC3^ and SCMC^ZBED3^ mediated by the LRRs of NLRP5. (C) Structural comparison of SCMC^KHDC3^ and SCMC^ZBED3^, illustrating the replacement of the NLRP4f PYD by ZBED3. (D) Dimerization of SCMC^KHDC3^ and SCMC^ZBED3^ in the adjacent bulbous units mediated by NLRP4f. In the upper unit, NLRP4f engages OOEP via a PYD-KH interaction, with KHDC3 contacting the OOEP C-terminal helix and the NLRP4f NACHT. In the lower unit, ZBED3 interacts with OOEP-KH, and NLRP4f LRR contacts ZBED3, NLRP5 LRR, and OOEP. (E-F) Volcano plots show the differentially expressed proteins (fold change > 1.5, *p*-value < 0.05) in *Ooep* KO (left) and *Tle6* KO (right) oocytes compared with WT. The down-regulated CPL components are marked in red. (G) Venn diagram shows the overlap of down-regulated proteins of *Ooep* and *Tle6* KO oocytes. The numbers in each section represent the count of differentially regulated proteins that are unique or shared between two KO oocyte samples. The common down-regulated CPL proteins are highlighted in red. (H) TEM images show the morphology of the CPL filaments in GV oocytes from *Ooep^+/−^* and *Ooep^⁻/⁻^* mice. Red arrowheads indicate CPL filaments.

To dissect the functional contribution of SCMC components to CPL assembly and oocyte function, we generated *Ooep^−/−^* mice (Figure S8A). Female *Ooep^−/−^* mice were infertile, and the majority of embryos arrested at the two-cell to four-cell stage (Figures S8B-S8D), consistent with previous reports^25,26^. Of note, all SCMC components were markedly reduced in abundance in *Ooep^−/−^*oocytes (Figures 3E-3G, S8E-S8I), mirroring observations in *Tle6* knockout mice^6^. This interdependence aligns with our structural finding that SCMC constituents assemble into a stable complex, wherein depletion of core members destabilizes the entire assembly. Ultrastructural analysis further demonstrated a complete absence of CPL filaments in *Ooep^−/−^* oocytes (Figure 3H). Indeed, depletion of any individual SCMC component has been reported to disrupt CPL formation, indicating that these proteins are collectively essential for CPL formation^27–30^ (Table S5).

### Ubiquitination-related proteins in CPLs

Within each asymmetric unit of the CPL bulbous repeat, two UHRF1-UBE2D3 heterodimers occupy equivalent binding sites on the PADI6 tubule, and each recruits one NLRP14 – the third NLRP family member identified in CPLs (Figures 4A-4C). This quasi-centrosymmetric geometry orients the UHRF1-UBE2D3 modules and their associated NLRP14 partners in opposing directions, placing one NLRP14 in a “closed” conformation that contacts its counterpart from the adjacent asymmetric unit, while the other NLRP14 adopts an “open” configuration (Figure 4A). Consistent with their direct physical interaction, loss of NLRP14 reduces UHRF1 protein abundance, and vice versa^31,32^.

**Figure 4.**
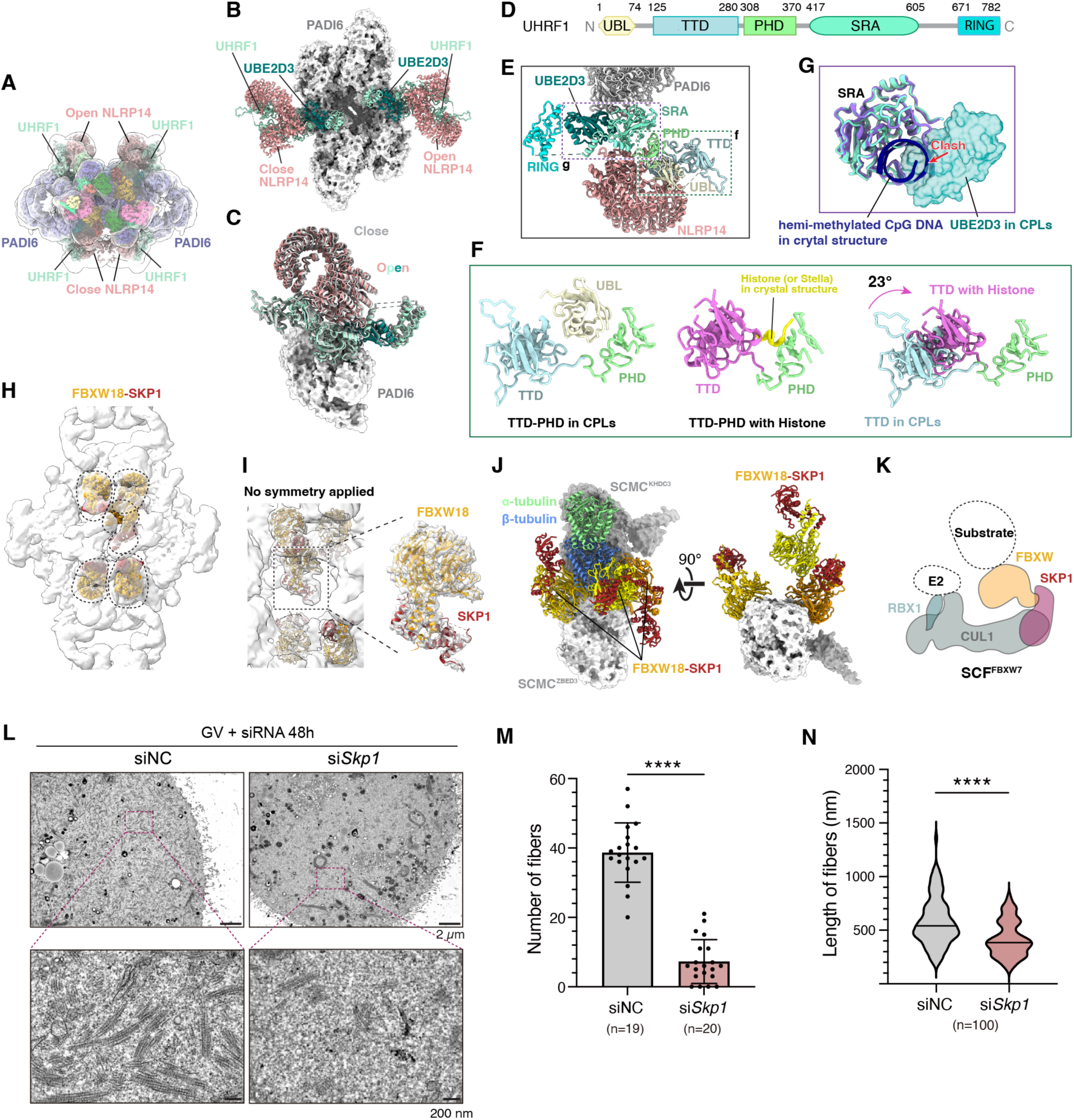
Structural organization of ubiquitination-related proteins within mouse CPLs. (A) Top-view cryo-EM map of CPLs showing NLRP14 in two conformations. Each protein component is shown with a distinct color. (B) Cryo-EM map of PADI6 tubule and its associated UHRF1-UBE2D3-NLRP14 modules. (C) Structural superposition of PADI6-UHRF1-UBE2D3-NLRP14 module in open and close conformations with the UHRF1-binding PADI6 as the reference. The r.m.s.d. is 2.8 Å across 1763 Cα pairs. (D) Schematic representation of the domain architecture of UHRF1. (E) Model showing UHRF1 domains bridging UBE2D3, PADI6, and NLRP14. (F) Structural comparison of the TTD and PHD of UHRF1 in the CPL and in complex with histone tail (PDB:3ASK). The TTD domains are colored in cyan and purple in the CPL and histone-bound complex, respectively. The TTD domain in the crystal structure is rotated by 23 degrees relative to the TTD domain in the CPL. (G) Structural comparison of the SRA of UHRF1 in the CPL and in complex with hemi-methylated CpG DNA (PDB:2ZKD). The SRA domains are colored in medium green and slate blue in the CPL and DNA-bound complex, respectively. The UBE2D3 in the CPL is shown as a surface. The hemi-methylated CpG DNA is shown in blue. (H) Cryo-EM map fitted with five SKP1-FBXW18 dimers within CPL. (I) The model of central SKP1-FBXW18 fitted with the cryo-EM map, reconstructed with C1 symmetry and low-pass filtered to 10 Å resolution (left). The enlarged view shows a well-resolved map reconstructed after symmetry expansion and local refinement (right). (J) Three SKP1-FBXW18 dimers associate with an SCMC dimer and α/β-tubulin heterodimer. (K) A cartoon diagram of SKP1-FBXW18 within the SFC complex based on EMDB: 14607. (L-N) *Skp1* knockdown reduces CPLs fiber number and length in mouse GV oocytes. Representative TEM images of GV oocytes 48 h after microinjection with siNC or si*Skp1* (L). Upper panels, 2,500×; lower panels, 10,000× magnification of the boxed regions. Quantification of CPLs fiber number from individual 10,000× TEM images (M; \*\*\*\**p* < 0.0001). Each dot represents one image; bars indicate mean ± s.d.. Violin plots of CPLs fiber length measured from 10,000× TEM images (N; \*\*\*\**p* < 0.0001); horizontal lines indicate medians.

UHRF1 is a multidomain RING-type E3 ubiquitin ligase with established non-canonical functions in DNA methylation homeostasis^33^. In CPLs, however, its domain organization adopts a configuration indicative of functional sequestration. The N-terminal ubiquitin-like (UBL) domain is inserted into the cavity formed between the tandem Tudor domain (TTD) and the plant homeodomain (PHD) (Figures 4D-4F). In comparison, prior crystallographic studies showed that in the absence of the UBL domain, this cavity narrows through a 23° hinge movement between TTD and PHD – a conformational state that accommodates histone tails or Stella to regulate DNA methylation in mammalian oocytes^34,35^. Occupation of this cavity by the UBL domain in the CPL structure would sterically preclude such interactions, rendering it inaccessible to its ligands (Figure 4F). UBE2D3, a broadly acting E2 enzyme that partners with multiple E3 ligases^36,37^ and is highly expressed in oocytes^38^, is tightly sandwiched between the central SET-and RING-associated (SRA) domain and the C-terminal RING domain of UHRF1 (Figure 4E). Remarkably, the SRA domain, which in nucleus binds hemi-methylated CpG DNA to direct epigenetic maintenance^39^, has its DNA-binding interface occluded by UBE2D3 within the CPLs (Figure 4G). UHRF1-UBE2D3 anchors to PADI6 primarily through the SRA domain (Figure 4E), consistent with the human PADI6-UHRF1-UBE2D architecture^40^. Collectively, these observations indicate that UHRF1 is maintained in an inactive conformation when incorporated into CPLs, with its key interaction surfaces for DNA, histones, and Stella sequestered by intra-complex contacts – a structural mechanism that likely preserves maternal factor quiescence until developmentally programmed release.

CPLs also harbor a second ubiquitin ligase-related module, a heterodimer of SKP1-FBXW18. Consistent with our structural assignments, FBXW18 was reproducibly detected in co-IP and mass spectrometry analyses targeting TLE6, OOEP and FLAG-KHDC3^41^. Given that multiple FBXW family members are highly expressed in mammalian oocytes and early embryos (Figure S4C), it is plausible that additional FBXW isotypes may be incorporated into CPLs as minor components. SKP1 and FBXW proteins are core components of SCF (SKP1-Cullin-F-box) E3 ubiquitin ligase complexes, which couple Cullin–RING proteins to specific substrates for proteasomal degradation^42^. Five SKP1-FBXW18 heterodimers were identified in each CPL repeating unit, with one SKP1-FBXW18 located asymmetrically at the central cavity (Figure 4H-I). In addition, two SKP1-FBXW18 dimers directly engage an SCMC dimer through the WD-repeat (WDR) domain of FBXW18 – the canonical substrate-recognition module of SCF complexes (Figures 4J and 4K). Notably, these SKP1-FBXW18 modules are deeply embedded within the CPL architecture, making them inaccessible to both the Cullin-RING catalytic machinery and potential substrates. This spatial sequestration suggests that FBXW18-SKP1 is stored within CPLs in an inactive state, thereby coordinating the temporal control of ubiquitin-mediated proteostasis during early development.

Expansion microscopy (ExM) has co-localized PADI6 with UHRF1, UBE2D3 and SKP1^6^. To determine how these enzymatic modules contribute to CPL assembly, we knocked down *Skp1* expression using siRNA in fully-grown mouse GV oocytes (Figure S9). *Skp1* depletion resulted in a significant reduction in both the number and length of CPL filaments relative to wild-type controls (Figures 4L-4N). Immunofluorescence analysis further revealed decreased SKP1 and UHRF1 signals, indicating that SKP1 is required for maintaining CPL integrity (Figures S9C and S9D). Similarly, siRNA depletion of *Ube2d3* in secondary follicles resulted in a pronounced loss of CPL filaments, accompanied by reduced UHRF1 levels (Figures S9E-S9J). Together, these results confirmed our structural assignments and demonstrated that the newly identified CPL components SKP1 and UBE2D3 are essential for CPL assembly.

### Tubulin heterodimers in CPLs

How do CPLs interact with the conventional cytoskeleton? We identified four α/β-tubulin dimers embedded within each bulbous unit of CPL filaments (Figure 5A)^43^. Within each asymmetric unit, one tubulin dimer associates with the open NLRP14, whereas the other engages the closed conformation, thereby positioning one α/β-tubulin dimer closer to the core of the bulbous unit and the SCMC, and the other at a more peripheral location (Figures 5B-5E). Despite these distinct spatial arrangements, the α-tubulin-NLRP14 interface is conserved between the two states: the E50-V57 β-strand of NLRP14 extends the S7-S10 β-sheet of α-tubulin to form a continuous β-sheet (Figure 5F). This interaction is further reinforced by the negatively charged C-terminal H12 helix of α-tubulin, which also contacts NLRP14, collectively enabling stable association (Figure 5G). In contrast, the β-tubulin exhibits differential interactions with SKP1-FBXW18, engaging through multiple structural elements including the β-strand S3, α-helixes H3, H11, H12, and S5-H5 and H11-H12 loops (Figures 5D, 5E and 5H). This multivalent binding capacity is consistent with the established role of SKP1-FBXW18 in recruiting diverse substrates for ubiquitination^42^.

**Figure 5.**
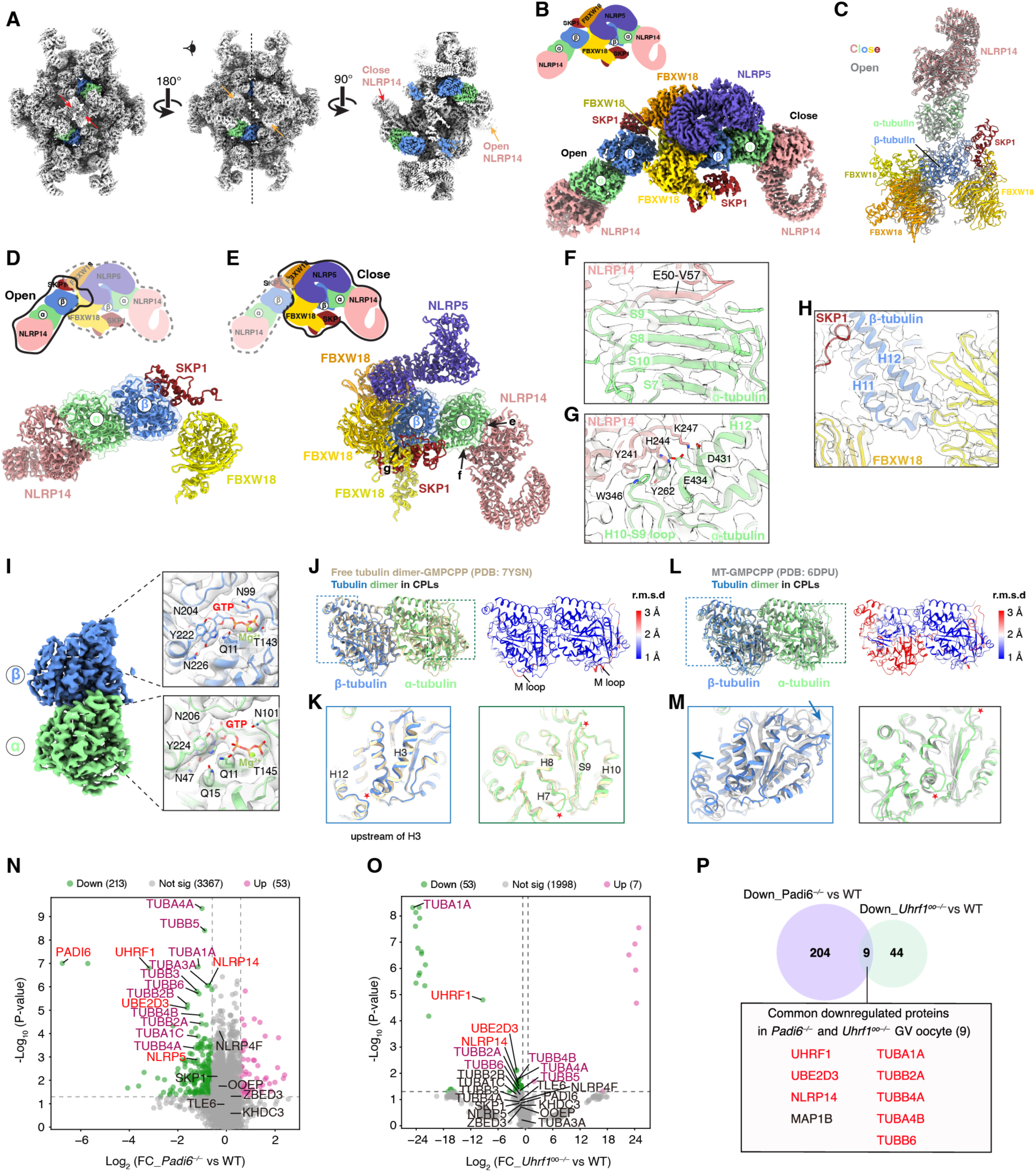
Structural organization of α/β-tubulin dimers within mouse CPLs. (A) Cryo-EM map of four α/β-tubulin heterodimers within one repeating unit. (B) Overall architecture of two α/β-tubulin heterodimers and their associated proteins. (C) Structural comparison of the α/β-tubulin heterodimers and their associated proteins in open and closed conformations, with two α/β-tubulin heterodimers superposed. (D-E) Structural model of the α/β-tubulin heterodimers and their associated proteins in open and closed conformations. (F-G) Interactions between α-tubulin and NLRP14 in the closed conformation. These interactions are similar in open and closed conformations of NLPR14. (H) β-tubulin interactions with SKP1 and two FBXW18 proteins in the closed conformations. (I) Cryo-EM map of the α/β-tubulin heterodimer showing densities of GTP and Mg²⁺ in both α-and β-tubulins. (J-K) Structural comparisons of the CPL-associated α/β-tubulin heterodimer with free α/β-tubulin (PDB: 7YSN). The r.m.s.d between the two structures is color-coded (J, right). Red stars mark the structural differences between the two structures in (K). (L-M) Structural comparisons of the CPL-associated α/β-tubulin heterodimer with MT α/β-tubulin (PDB: 6DPU). The r.m.s.d between the two structures is color-coded (L, right). Red stars and blue arrows mark the structural differences between the two structures (M). (N-O) Volcano plots show the differentially expressed proteins (fold change > 1.5, *p*-value < 0.05) in *Padi6* KO (left) and *Uhrf1* KO (right) oocytes compared with WT. Tubulins are marked with purple and other CPL components are marked with red. (P) Venn diagram shows the overlap of down-regulated proteins of *Padi6* KO and *Uhrf1* KO oocytes. The numbers in each section represent the count of differentially regulated proteins that are unique or shared between two KO oocytes samples. Tubulins and other CPL components are highlighted in red.

The high-resolution cryo-EM map enables unambiguous assignment of the nucleotide states, revealing that both α– and β-tubulins are bound to GTP and Mg²⁺ (Figure 5I). Structural superposition further shows that tubulins within CPLs closely resemble the conformation of GMPCPP-bound tubulin dimers, but differ markedly from the tubulin conformation observed in assembled microtubules (MTs) (Figures 5I-5M). Beyond the intrinsically flexible M loop, the most prominent deviations between CPL-bound and free tubulins are localized primarily to the upstream region of the β-tubulin H3 helix and the α-tubulin H7-H8 and S9-H10 loops, all of which lie proximal to interfaces with other CPL components (Figure 5K). These subtle differences indicate that incorporation of α/β-tubulin dimers into CPLs requires only minimal structural remodeling. The retention of a GTP-bound, assembly-competent conformation suggests that CPLs harbor a reservoir of polymerization-ready tubulins, poised to support rapid MT assembly upon release.

Tubulins are abundantly expressed in oocytes and early embryos (Figure S4B). Reanalysis of published proteomic data from *Padi6* or *Uhrf1* knockout mouse oocytes revealed a marked reduction in both α– and β-tubulins, accompanied by decreased levels of UHRF1, UBE2D3 and NLRP14 (Figures 5N-5P). In contrast, tubulin abundance is not significantly altered in *Ooep* or *Tle6* knockout oocytes (Figures 3E and 3F), despite the loss of CPL filaments. The maintenance of tubulin levels in the absence of intact CPL filaments or SCMCs suggests that these CPL factors, other than SCMCs, can stabilize tubulin independently of filament assemblies, likely by forming intermediate subcomplexes. Together, these findings support a model in which CPL proteins not only scaffold filament architecture but also act as molecular stabilizers that preserve the oocyte tubulin reservoir under conditions of structural perturbation.

### High-order assembly of CPLs

The parallel arrangement of CPL filaments observed in cryo-EM and TEM images prompted us to investigate whether CPLs assemble into higher-order architectures (Figure 1A and 4L). By increasing the particle box size and performing focused 3D classification, we resolved four pairs of double-filament CPLs which could be further assembled into a complete five-helical bundle structure (Figures 6A and S1G). Notably, CPL units adopt a tilted packing along the filament axis, with an inter-unit tilt of 6-7 degrees (Figure 6B), which is consistent with the cryo-ET observation^6^, further validating that our CLoG sample-preparation strategy preserves CPLs in a near-native state.

**Figure 6.**
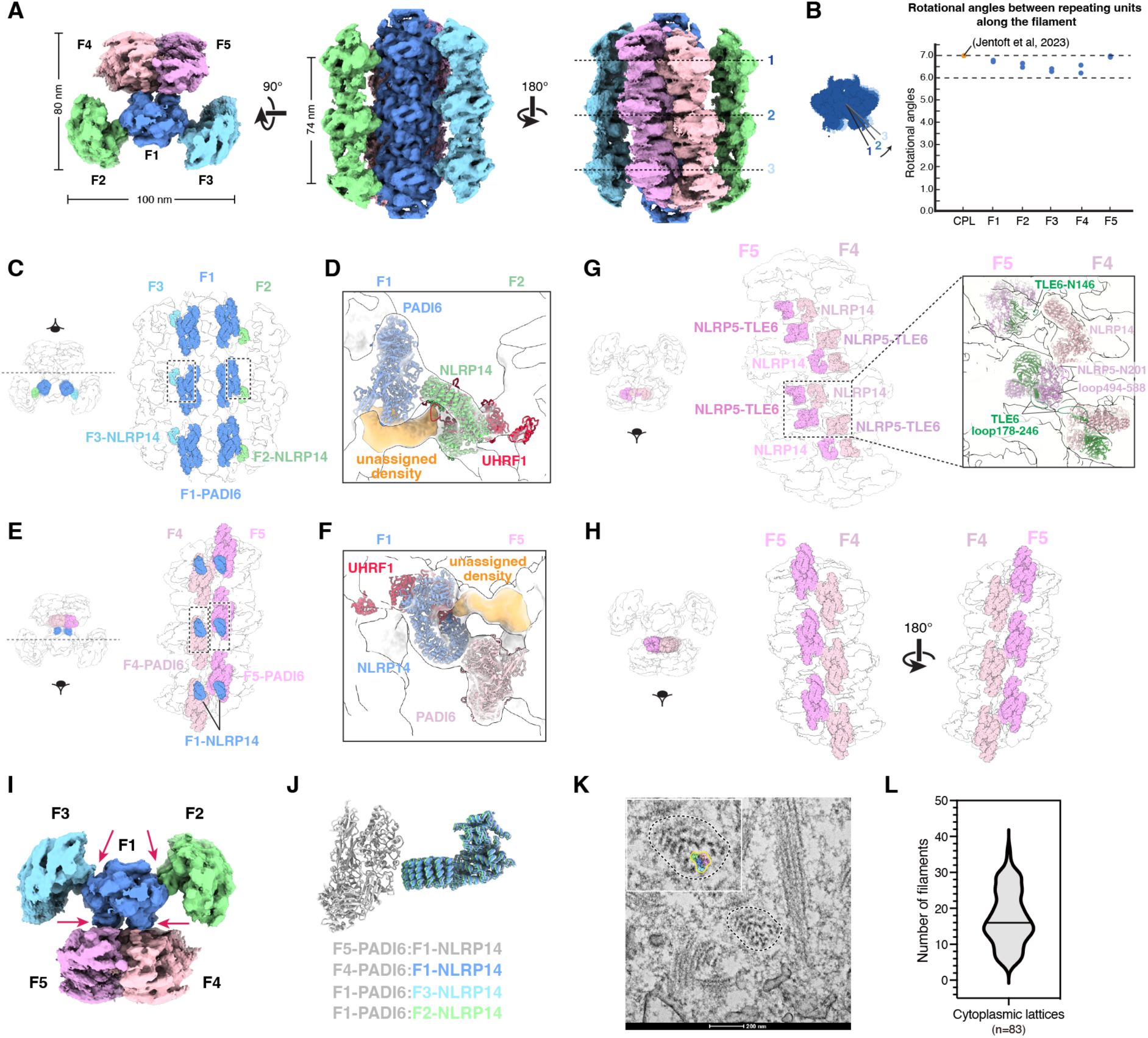
High-order assembly of CPL bundles. (A) Cryo-EM map of the five-helical CPL bundle shown in side and end views, with individual filaments (F1–F5) coloured differently. Dashed lines indicate repeating units along the filament axis that were used as markers to calculate the rotational angles in (B). (B) Rotational angles between adjacent repeating units along individual CPL filaments, compared with previously reported cytoplasmic lattice parameters. (C) Interactions between the central filament F1 and external filaments F2 and F3 mediated by NLRP14 and PADI6. (D) Zoom-in view of the boxed region in c, illustrating the contacts between F1-PADI6 and F2-NLRP14 and UHRF1, with the UHRF1 loop (residues 76-125) indicated. (E) Interactions between F1 and filaments F4 and F5 mediated by NLRP14 and PADI6. (F) Zoom-in view of the boxed region in e, illustrating the contacts between F1-NLRP14 and UHRF1 loop (residues 76-125) and F4-PADI6. (G) Interactions between filament F4 and F5 mediated by NLRP5-TLE6. (H) Zoom-in view of the boxed region in (G), illustrating interactions between NLRP5-TLE6. (I) Interactions between filament F4 and F5 mediated by PADI6. (J) Interactions between filaments mediated by NLRP14 and PADI6. Superposition of NLRP14 and PADI6 at different filament interfaces, with PADI6 as the reference. (K) Representative TEM micrographs of CPL filaments observed in oocytes, with dashed outlines indicating filamentous assemblies corresponding to CPL bundles. (L) Violin plot showing the distribution of filament numbers within CPL bundle assemblies (n = 83). Horizontal lines indicate medians.

Within this assembly, four peripheral filaments (F2-F5) associate with a central filament (F1) through conserved NLRP14-PADI6 interactions (Figures 6C-6F). In addition to NLRP14-mediated contacts, UHRF1 likely contributes to inter-filament coupling by engaging an unassigned density of neighboring filaments (Figures 6D and 6F). Further multivalent interactions involving PADI6, open-state NLRP14 and SCMC components from the F4 and F5 filaments reinforce lateral association (Figures 6G and 6H). Together, these findings establish that open-state NLRP14, UHRF1 and PADI6 – which are exposed on the outer surface of the bulbous unit – serve as principal mediators of inter-filament connectivity (Figures 6I and 6J). Notably, the five-filament bundle likely represents a fundamental assembly intermediate, as each CPL fiber in mouse oocytes contains, on average, more than 15 filaments, indicating progressive lateral accretion into higher-order supramolecular structures (Figure 6K and 6L).

### CPL proteins are associated with human variants

Many of the CPL components are maternal-effect genes and variants in these genes have frequently been linked to female infertility, manifesting as early embryonic arrest, multilocus imprinting disorders or hydatidiform molar pregnancies^22,44^ (Figure S10, Tables S6 and S7). SCMC components (*NLRP5, TLE6, OOEP, KHDC3L*), *PADI6* and *NLRP14* have been implicated in early embryonic arrest^9–12^, with *NLRP5, KHDC3L and PADI6* additionally associated with hydatidiform moles^45–47^. Furthermore, variants in *UHRF1,* as well as in *NLRP5*, *OOEP* and *PADI6,* have been shown to cause multilocus imprinting disturbances^48,49^ (Table S6). Of note, while variants in the β-tubulin *TUBB8* and the α-tubulins *TUBA4A* and *TUBA1C* are known to cause early embryonic arrest or oocyte maturation abnormality in humans^50–52^, their involvement in CPL assembly has not yet been demonstrated.

We mapped missense mutations onto the CPL structural model (Figure S10C, Table S7), including variants in *NLRP5* (31), *TLE6* (15), *OOEP* (1), *PADI6* (31), and *UHRF1* (1). Several mutations are located at surface-exposed sites, such as OOEP R32; TLE6 T305, R417 and E276; NLRP5 R699 and L1079; and PADI6 N586, H425, D337 and R340, where they may participate in protein-protein interactions. In contrast, most mutations are buried within the protein interior and may either affect hydrophobic core formation (e.g., NLRP5 F554 and L602) or disrupt intramolecular interactions, potentially compromising structural stability. Notably, many recurrent embryo developmental problems associated with CPLs cannot be overcome by intracytoplasmic sperm injection (ICSI)^53^, highlighting the importance of understanding the genetic basis of CPL-related female infertility and the value of genetic diagnosis prior to pregnancy.

## Discussion

By developing an efficient CLoG cryo-EM sample preparation strategy, we determined the near-atomic structure of endogenous CPL filaments, enabling *de novo* building of their atomic models. We identified fourteen protein components within the CPL – comprising both well-characterized and previously understudied factors – and elucidated their detailed molecular interactions. These findings provide a mechanistic foundation for understanding how oocyte proteins govern CPL assembly and early embryonic development.

### SCMC and PADI6 form the core CPL scaffold

The reconstituted SCMC core (NLRP5, TLE6 and OOEP) forms a stable complex that can dimerize^13^. Our structure further reveals that incorporation of accessory SCMC proteins (KHDC3 and ZBED3) together with NLRP4f organizes four SCMCs into a hollow, super-tetrameric assembly. Depletion of any of these five SCMC proteins or NLRP4f disrupts CPL formation in mouse (Table S6). Notably, loss of KHDC3, ZBED3, or NLRP4f does not markedly affect expression of the SCMC core proteins, whereas depletion of TLE6 or OOEP drastically reduces the levels of all SCMC components^29,30,54^ (Figures 3E and 3F). Consistently, knockout of *Khdc3*, *Zbed3*, or *Nlrp4f* results in delayed preimplantation development and reduced fecundity – phenotypes less severe than the early embryonic arrest and female sterility observed in *Nlrp5*, *Tle6*, or *Ooep* knockout mice^26,29,30,54,55^ (Figure S10A, Table S6). This asymmetric dependency indicates that while accessory proteins are required for proper CPL assembly, the SCMC core subcomplex remains intact in their absence and can partially support oocyte function.

Another unexpected finding is that PADI6 dimers self-assemble into hollow tubules both in native CPLs and *in vitro*, revealing an inherent capacity for oligomerization. These PADI6 tubules interconnect with super-SCMC tetramers throughout the CPL, together forming its principal structural scaffold. Interestingly, most SCMC components were not particularly enriched by Flag-PADI6 pull-down (Figure S5B), and their protein levels remained largely unchanged upon *Padi6* knockout (Figure 5N). These observations suggest that PADI6 and the SCMC initially assemble into distinct subcomplexes that are subsequently integrated into CPL filaments through relatively weak interactions. In contrast, the abundance of the UHRF1-UBE2D3 complex and NLRP14 is strongly dependent on PADI6, and this dependency is unidirectional, as depletion of UHRF1 does not affect PADI6 levels (Figures 5N-5P). This asymmetric dependency mirrors the asymmetric reliance of accessory SCMC components on the SCMC core, further supporting a modular-centered model of CPL assembly.

### CPLs sequester ubiquitination and cytoskeletal machineries

Remarkably, ubiquitination modules and cytoskeletal α/β-tubulin are positioned at the center of each CPL bulbous unit. Functional ablation of *Uhrf1* completely abolishes CPL formation^32^, while knockdown of *Ube2d3* or *Skp1* markedly reduces CPL abundance, indicating that these modules are integral to CPL assembly. Although the filamentous organization of CPLs resembles enzyme polymers observed in eukaryotic cells (e.g., CTP synthase and acetyl-CoA synthetase)^56,57^, CPLs are fundamentally distinct in integrating more than a dozen proteins that span diverse cellular pathways, including protein degradation, DNA methylation, cytoskeletal dynamics, and DNA repair.

The presence of ubiquitination machinery further suggests that CPLs function either as a regulated degradation platform or as a storage depot. Our structures reveal that UHRF1 is maintained in an autoinhibited state, with its ligand-binding sites occupied by its UBL domain and its DNA-binding domain occluded by the associated E2 ligase UBE2D3. Similarly, the SKP1-FBXW18 module is sequestered in a conformation inaccessible to Cullin-RING E3 ligases. Thus, these ubiquitination complexes are likely rendered inactive within CPLs through dual mechanisms: active-site blockade and sequestration within a fibrous array. Upon CPL disassembly during early embryogenesis, these previously restrained degradation factors are likely released and reactivated, enabling timely clearance of maternal proteins that are beneficial during oocyte maturation but detrimental to subsequent development^58^.

### Insights into CPL assembly

The structural organization of CPLs and the identification of subcomplexes, together with the unidirectional dependencies between the SCMC core and accessory factors, and between PADI6 and its partners, support a hierarchical assembly pathway for CPL filaments (Figure 7). Lateral interactions involving NLRP14, UHRF1 and PADI6 further promote inter-filament associations, giving rise to the characteristic five-helical-bundle configuration.

**Figure 7.**
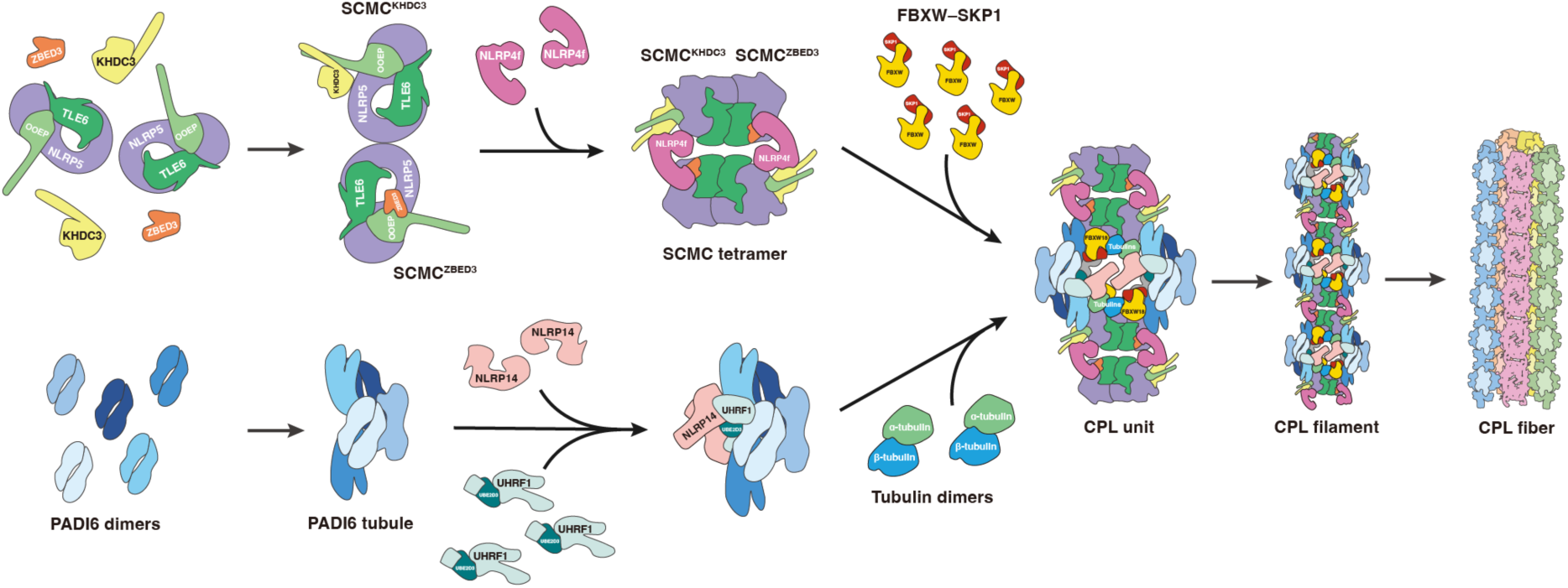
A hierarchical assembly model of CPLs. (i) SCMC core recruits accessory proteins KHDC3 and ZBED3 to assemble into two distinct SCMC complexes, which are bridged by NLRP4f to form the tetrameric SCMC. (ii) PADI6 dimers assemble into PADI6 tubule, which recruit UHRF1, UBE2D3 and NLRP14. (iii) Tubulin dimer and SKP1-FBXW18 modules are incorporated with SCMCs and PADI6 tubules to form the CPL filament. (iv) Five CPL filaments assemble into a helical bundle through inter-filament interactions mediated by NLRP14 and PADI6, which recruit more CPL filaments to form the lattice.

In addition to the core components defined here, numerous proteins – including SPIN1, YWHA family members and PARP1 – have been reported to associate with CPLs^6,41^ (Table S4). Our structure indicates that these factors are not integral components of CPLs but are likely transient or low-occupancy interactors. The extensive exposed surfaces and unresolved flexible regions within CPL proteins provide potential docking platforms for such factors (Figure S3B). Consistent with this idea, XL-MS identified a linkage between the unresolved C-terminal tail of OOEP (K147) and PARP1 (K134) (Table S5). Our structural analysis further excludes ribosomes and intermediate filaments as intrinsic components of CPLs^17,24,25,48,49^. Nevertheless, ribosomal proteins are enriched in PADI6 pull-down assays (Figure S5), suggesting that ribosomes or their components may associate with CPLs through peripheral or dynamic interactions.

What is the mechanism underlying CPL disassembly? NLRPs comprise a family of more than ten proteins in mammals with established roles in immune signaling and oocyte function. Studies of NLRP3, a well-characterized inflammasome component, have shown that ATP hydrolysis triggers dramatic conformational rearrangements essential for its activation^61,62^. While the recombinant NLRP5 in SCMC shows no ATP density in its conserved binding pocket^13,14^, all three NLRP proteins exhibit bulky densities in corresponding regions of our mouse CPL structure, suggestive of bound ligands, although the molecular nature of these ligands remains ambiguous due to resolution limitation (Figure S11). This raises the possibility that ATP hydrolysis-driven conformational changes in CPL-associated NLRPs may initiate filament disassembly – a hypothesis warranting future investigation.

Our study revealed the core components and hierarchical assembly of mouse oocyte CPLs at unprecedented resolution. By presenting functional protein modules in an inactive, polymerized state, the CPL architecture may serve as a sequestration depot, enabling the timely release and deployment of factors for post-fertilization processes such as protein degradation and cytoskeletal remodeling. Our work also provides a framework for interpreting pathogenic variants in CPL genes linked to female infertility, offers a refined gene panel to enhance molecular diagnostics, and lays the foundation for targeted therapeutic strategies aimed at addressing fertility disorders and genomic imprinting defects.

## Methods

### Animals

Wild-type (WT) female ICR mice of 3-4 weeks old were used for the preparation of the CPL sample in this study, which were obtained from the Zhejiang Academy of Medical Science, China. Mice maintenance and experiments were conducted in accordance with the institutional guidelines and approved by the Institutional Animal Care and Use Committee of Zhejiang University.

### Oocyte collection and embryo culture

Female mice of 3-4 weeks old were injected intraperitoneally with 5 IU pregnant mare serum gonadotrophin (PMSG; Sansheng, 110254564) and were humanely euthanized with CO_2_ asphyxiation 44-48 hours later. Ovaries were mechanically cut into pieces, and GV stage oocytes were harvested and cultured in M2 medium (Sigma-Aldricn, M7292) with 2.5μM milrinone (MCE, HY14252) covered with mineral oil (Sigma-Aldrich, M5310) at 37 °C in a 5% CO_2_ atmosphere before use. Zygotes were harvested from fertilized females 24 h after the hCG injection and cultured in KSOM medium (Aibei, M1435).

### CPL sample preparation for negative staining

Carbon-coated copper grids were glow-discharged (CoolGlow; 25 W, 60 s, 40 Pa) immediately before use. GV-stage oocytes were resuspended in incubation buffer and mechanically disrupted by 1,000 strokes using a pestle. The lysate was clarified by centrifugation at 500g, after which virtually no intact oocytes remained, and the supernatant containing CPLs was collected for negative-staining sample preparation. A 3-μL aliquot of sample was applied to the carbon surface, incubated for 1 min, and blotted with filter paper. The grid was washed once with 3 μL of staining solution and blotted. For CPLs, a second 3-μL application of staining solution was incubated for 30 s, whereas for purified proteins, this step was incubated for 1 min. Prepared grids were transferred to labeled storage boxes and imaged on a Talos L120C transmission electron microscope (Thermo Fisher Scientific).

### CPLs sample preparation for cryo-EM

Based on the previous description, fully grown oocytes (>70 µm) at the germinal vesicle (GV) stage were harvested. The incubation buffer was prepared according to previously published protocols^63^, containing 10 mM PIPES buffer (pH=7.2), 100 mM NaCl, 3 mM MgCl₂,and 10 mM EGTA, supplemented with 1x cOmpleteTM EDTA-free Protease Inhibitor Cocktail [Roche, RF(m)11647] and 1 mM phenylmethylsulfonyl fluoride (PMSF; Solarbio, IP0280). Collected GV oocytes were washed three times in fresh incubation buffer.

An aliquot of 2.5 μL of the oocyte suspension (about 400-500 oocytes) was applied onto a glow-discharged Quantifoil Au R2/1, 300-mesh grid coated with a continuous carbon film and incubated for 1 min. To permeabilize the oocytes, 0.5 μL of lauryl maltose neopentyl glycol (LMNG; Anatrace, NG310) solution, prepared as a 6× detergent stock in distilled water, was added and gently mixed to achieve a final concentration of 0.05%. The oocytes were then incubated at room temperature for 10 min, with the incubation buffer replenished as necessary to prevent drying. The grids were blotted for 2–6 s and rapidly plunge-frozen in liquid ethane cooled by liquid nitrogen using a Vitrobot Mark IV (Thermo Fisher Scientific) operated at 100% humidity and 4 °C.

### Cryo-EM data collection of mouse CPLs

Cryo-EM data of the CPLs was collected in four sessions in the cryo-EM facility of Liangzhu Laboratory or ShuimuBio at a nominal magnification of 130, 000 × on a Titan Krios G4 (Thermo Fisher Scientific) operating at 300 kV equipped with a Falcon 4i detector and Selectris X energy filter (slit width 20 eV). Movie stacks were automatically acquired using EPU (Thermo Fisher Scientific). The defocus range was set from –1.5 ∼ –2.5 µm and the total dose per movie stack was ∼ 50 e^−^/Å^2^.

### Image processing of mouse CPLs

The data processing workflow was summarized in Figure S1. Microscope settings are summarized in Table S1. Initial particle selection was performed separately for each session. Movie stacks were motion corrected using RELION’s implementation^64^ and CTF estimation was performed with CTFFIND 4.1^65^. Particle picking was the most challenging and time-consuming part of the data processing as all mouse oocyte proteins are in the images, resulting in a low contrast. We utilized multiple particle picking tools including CryoSPARC filament tracer^66^, Topaz^67^, CrYOLO^68^ and manual picking in RELION5. For artificial-intelligence (AI) powered software like Topaz and CrYOLO, multiple rounds of training were performed using the selected particles to improve the AI model. Particles were extracted with 18-nm spacing along the filaments, using a box size of 640, which were down-scaled 4 times to accelerate image processing. Extensive rounds of 2D classification were performed to select good particles. Particles selected from different tools were joined and duplicates were removed. Subsequent data processing was performed in RELION unless otherwise stated. 3D refinements were performed using the cryo-ET density map of mouse CPLs (EMD-16458) as the initial model. A longitudinal periodicity of ∼ 38 nm and a C2 symmetry were identified. To recover more particles, each particle’s coordinate was shifted ± 38 nm along the longitudinal axis, to its adjacent units. The expanded particle set was joined and duplicates were removed. After rounds of 3D refinements and 3D classifications, good particles were selected for this session. All the datasets from 4 sessions were processed similarly and good particles were joined, yielding 288,673 particles. Selected particles were re-extracted with bin 1, followed by 3D classification, CTF refinement and Bayesian polishing. 3D refinement with Blush regularisation got a 4.8 Å density map with C2 symmetry. Then the particle symmetry was expanded to C1 and one asymmetric unit was subtracted. The boxsize of particles corresponding to half of the CPL unit was reduced to 400, yielding a 4.3 Å density map. 3D classification and refinements with local masks improved the densities up to 3.4 Å.

To improve the narrow segment, the selected good particles from the 4 datasets were shifted ± 19 nm, along the longitudinal axis. After 3D classification, CTF refinement, Bayesian polishing and 3D refinement, a consensus density map with C2 symmetry was obtained with a resolution of 4.4 Å. Symmetry expansion, subtraction, and boxsize reduction were performed to obtain an improved density map of the connecting region between CPL units. The density around NLRP14 was refined to 4.0 Å. The local refined density maps were stitched together in UCSF ChimeraX^69^ using *volume maximum* to generate a composite map of an asymmetric unit CPL.

To recover the densities of adjacent filaments, particles were re-extracted with a boxsize of 1280 and down-scaled 4 times. 4 filaments can be observed around the central CPL filament. 3D classifications were performed to recover the density of adjacent filaments, each time focused on one filament. Four pairs of CPL filaments were obtained by 3D refinements. These filaments were stitched together in UCSF ChimeraX to generate the five-helical-bundle fiber. The map resolution was calculated based on the Fourier shell correlation (FSC) = 0.143 criterion. Local resolutions were calculated using RELION5

### Model building and refinements

Strategies for the identification of CPL proteins were summarized in Supplementary Table 2. Protein identification was performed by an integrated method combining AI-guided structural predictions using Alphafold2^70^, Alphafold3^71^ and AlphaPulldown^72^, automatic fitting of structural predictions into density maps by Colores in Situs^73^, automatic fold recognition by FoldSeek^74^, AI-based *de novo* modeling by ModelAngelo^75^, fitting of known structural fragments and manual inspection. Protein assignments were confirmed by sidechain densities. The isotypes of ɑ-tubulin and β-tubulin were uncertain and were tentatively assigned as TUBA1c (P68373) and TUBB2a (Q7TMM9). based on their relative abundancy in mass spectrometry analyses. Models were fitted into cryo-EM density in UCSF ChimeraX and manually refined in Coot^76^. The final atomic model was refined against its corresponding density map using Phenix^77^ with secondary structure, Ramachandran and rotamer restraints applied. Iterative correction with Coot was performed between rounds of Phenix refinement. Model statistics were generated using phenix.molprobity and are provided in Table S1.

### Preparation of CPLs and cross-linking for mass spectrometry

For cross-linking and mass spectrometry analysis, GV-stage oocytes were isolated aseptically and suspended in incubation buffer containing 0.05% LMNG (final concentration, from a 0.5% stock solution). The samples were mechanically homogenized with a pestle for approximately 1 min (about 100 strokes), until most GV structures were disrupted as observed. Bis(sulfosuccinimidyl) suberate (BS3; Thermo Fisher Scientific, 21580) was added to a final concentration of 2 mM, and the mixture was incubated on ice for 30 min to allow cross-linking. The reaction was quenched by adding Tris buffer to a final concentration of 50 mM and incubating at room temperature for 15 min.

The samples were centrifuged at 3,000 × g for 10 min at 4 °C, and the supernatant was collected. The remaining material was further centrifuged at 20,000 × g for 10 min at 4 °C. The resulting pellet – referred to as CPLs – was retained and verified by negative staining. The CPLs pellet was resuspended in 30 μL of buffer, mixed with 5× Protien Sample Loading Buffer (Beyotime, P0286), and boiled at 96 °C for 8 min. The CPLs samples were subjected to Bis-Tris PAGE (GenScript, M00656) for approximately 5 min until all samples had completely entered the resolving gel. The gel was then fixed and stained using Coomassie Brilliant Blue staining solution (Beyotime, P0017). The corresponding protein bands and gel wells were excised and transferred into double-distilled water (ddH₂O) for liquid chromatography – tandem mass spectrometry (LC-MS/MS) analysis.

### LC-MS/MS Analysis of cross-linked CPLs

Gel bands corresponding to proteins exhibiting catalase activity were excised for in-gel digestion, and proteins were identified by MS as previously described^78^. Briefly, the proteins were reduced with 5 mM dithiothreitol (DTT; Biosynth, D-8200) and alkylated with 11 mM iodoacetamide. In-gel digestion was performed using sequencing-grade modified trypsin in 50 mM ammonium bicarbonate at 37 °C overnight. Peptides were extracted twice with 1% trifluoroacetic acid (TFA) in 50% acetonitrile aqueous solution for 1 h each. The combined peptide extracts were concentrated using a SpeedVac concentrator to reduce the volume. The resulting peptides were resuspended in 20 μL of 0.1% TFA and centrifuged at 20,000 × g for 15 min at 4 °C to remove particulate impurities.

For LC-MS/MS analysis, peptides were separated on a Vanquish Core liquid chromatography system (Thermo Fisher Scientific) coupled directly to a Thermo Orbitrap Exploris 480 mass spectrometer. Separation was performed at a flow rate of 0.3 μL/min using a 40 min’ linear gradient on a homemade fused-silica capillary column (75 μm ID, 150 mm length; Upchurch, Oak Harbor, WA) packed with C18 resin (300 Å, 5 μm; Varian, Lexington, MA). Mobile phase A consisted of 0.1% formic acid in water, and mobile phase B consisted of 100% acetonitrile with 0.1% formic acid.

The Orbitrap mass spectrometer was operated in data-dependent acquisition (DDA) mode using Xcalibur 4.0.27.10 software. Full MS scans were acquired in the Orbitrap over an m/z range of 300-1500 at a resolution of 120,000, followed by data-dependent MS/MS scans for 3 s in the Ion Routing Multipole using higher-energy collisional dissociation (HCD) with a normalized collision energy of 40%.

Cross-linking data were analyzed using pLink2 software. The following search parameters were applied: MS1 tolerance = ± 20 ppm; MS2 tolerance = ± 20 ppm; enzyme = trypsin (full tryptic specificity, ≤ 3 missed cleavages allowed); cross-linker = SDA (with lysine as one of the cross-linked residues); fixed modification = carbamidomethylation on cysteine; variable modifications = oxidation on methionine and acetylation at the N-terminus. A false discovery rate (FDR) < 5% was applied. α-fragments resulting from additional cleavage of cross-linked peptides at the lysine isopeptide bond were manually validated. The cross-linking data was visualized using the xiVIEW web application.

### Protein expression and purification

The full-length cDNAs of mouse wild-type (WT) PADI6, mutant PADI6 (R43A and W66), and WT PADI1 were separately cloned into a pCMV vector with an N-terminal Flag– and Strep-tag. Recombinant proteins were expressed using the HEK293F cell expression system. The cells were cultured at a density of 2 × 10⁶ cells/mL. For each 1 L culture, 2 mg of plasmid DNA was used for transfection. Polyethyleneimine (PEI; Polysciences, 23966) was added at a DNA-to-PEI mass ratio of 1:3. Cells were harvested 48 h post-transfection by centrifugation at 1,500 × g for 10 min, and the supernatant was discarded. The cell pellet from each liter of culture was resuspended in 50 mL of lysis buffer containing 20 mM Tris-HCl (pH 7.5), 200 mM NaCl, 10% (v/v) glycerol, 1 mM PMSF, 1 mM DTT, and a protease inhibitor cocktail. The cells were lysed by ultrasonication, and the lysate was clarified by centrifugation at 18,000 × g for 30 min at 4 °C. The supernatant was incubated with pre-equilibrated anti-Flag G1 affinity resin (GenScript, L00432) on a rotator at 4 °C for 30 min. After incubation, the resin was washed twice with lysis buffer, and bound proteins were eluted with 3×Flag peptide solution (0.1 mg/mL) prepared in lysis buffer. The eluate was concentrated and further purified by size-exclusion chromatography (SEC) on a Superdex 200 Increase 10/300 GL column (Cytiva, 28990944) equilibrated with lysis buffer. Protein-containing fractions were pooled, concentrated, and subsequently loaded onto a Capto HiRes Q 5/50 ion-exchange column (Cytiva, 29275878) equilibrated with 20 mM Tris-HCl (pH 7.5) and 50 mM NaCl. The proteins were eluted using a linear NaCl gradient. The peak fractions were collected, concentrated and aliquoted for further use. During the concentration step, the buffer was replaced with a glycerol-free lysis buffer to ensure sample compatibility with subsequent structural analyses.

### Gradient preparation and GraFix

WT and mutant PADI6 protein polymers were prepared using a modified GraFix protocol as previously described^23^. Briefly, a continuous sucrose–glutaraldehyde gradient was formed ranging from 10% to 25% (w/v) sucrose and 0.01% to 0.1% (v/v) glutaraldehyde (GTH; Sigma-Aldrich, 49626). A total of 0.3 mg of freshly puried PADI6 protein in buffer containing 20 mM HEPES (pH 7.5), 200 mM NaCl, 1 mM PMSF, 1 mM DTT, and a protease inhibitor cocktail was gently layered onto the top of the gradient and subjected to ultracentrifugation at 240,000 × g for 18 h at 4 °C (maximum acceleration, minimum deceleration). After centrifugation, a white pellet was visible at the bottom of the tube. The gradient was fractionated from top to bottom into 14 fractions, and the cross-linking reaction was quenched by adding Tris buffer to a final concentration of 20 mM.

SDS-PAGE analysis revealed thet PADI6 protein bands first appeared in fraction 4. Fractions 4–14 were pooled and concentrated separately, followed by buffer exchange to remove sucrose. The resulting sample was examined by negative staining to assess particle integrity and homogeneity. Fractions 9–12 were collected, pooled together, and concentrated to the minimal volume prior to cryo-EM grid preparation.

### Preparation of PADI6 protein samples for cryo-EM

To prepare cryo-EM grids, 3.5 μL aliquots of freshly purified WT PADI6 were vitrified under two conditions: (1) at a concentration of 0.3 mg/mL on glow-discharged Quantifoil Au R1.2/1.3, 400-mesh grids, and (2) at a concentration of 2 mg/mL supplemented with 0.01% LMNG on glow-discharged Quantifoil Cu R1.2/1.3, 400-mesh grids. The GraFix-treated PADI6 samples were vitrified on glow-discharged Quantifoil Au R1.2/1.3, 400-mesh grids. The grids were blotted for 3–5 s and rapidly plunge-frozen in liquid ethane cooled by liquid nitrogen using a Vitrobot Mark IV operated at 100% humidity and 4 °C.

### Cryo-EM data collection and data processing of PADI6

Cryo-EM data of the PADI6 untreated sample, the LMNG high concentration sample and the GraFix sample were collected using the same parameters as the CPL sample, and were processed using RELION 5.0 and CryoSPARC 4.5.1. The overall data processing workflow was summarized in Extended Data Figure 6. Movie stacks were initially pre-processed using RELION’s implementation for motion correction and CTF estimation with CTFFIND 4.1. Following pre-processing, particle picking for the untreated dataset was performed using Gautomatch with particle diameters of 150, 200, 250, 300 Å. Particle picking for LMNG high concentration dataset was carried out using both Gautomatch with particle diameters of 150, 200, 250, and 300 Å, and the CryoSPARC filament tracer with filament diameters of 60 Å and 110 Å. Particles were extracted with a binning factor of 4 and subjected to 2D classification. Particles selected from good-quality 2D class averages of the untreated dataset were used for 3D classification with C2 symmetry applied, using an initial model generated from the human PADI6 dimer. Particles from the good class were subsequently subjected to 3D refinement and additional rounds of 3D classification, together with PADI6 dimer particles selected from 2D classification of the LMNG high concentration dataset. Multiple rounds of 3D classifications were performed with C2 symmetry imposed. In the final round of 3D classification, C2 symmetry was applied and multi-references generated from a 3.8 Å cryo-EM map low-pass filtered to 3.8, 15, 25, and 35 Å were used. Particles with the highest resolution were selected from multi-reference 3D classification and further subjected to CTF refinement and polishing, yielding a final reconstruction map at an overall resolution of 3.0 Å.

Filamentous particles of 2D class averages from the LMNG high concentration dataset were removed duplicates within a distance threshold of 250 Å, re-extracted with a binning factor of 2, and subjected to non-uniform refinement using a 5-dimer cryo-EM map from CPL as the initial model, resulting in a reconstruction at an overall resolution of 9.1 Å.

Particle picking for the GraFix dataset was performed using a combination of multiple approaches to comprehensively identify particles, including Gautomatch with particle diameters of 150, 200, 250, and 300 Å; CryoSPARC filament tracer with filament diameters of 60 Å and 110 Å; CryoSPARC filament tracer using a template generated from the 5-dimer PADI map of CPL. All particles were extracted with a binning factor of 4 and subjected to multiple rounds of 2D classification. Particles with high-quality 2D class averages were selected. Duplicates were removed within a distance threshold of 250 Å, and subjected to further 3D classification. The first round of 3D classifications was performed using RELION 3D classification with alignment and CryoSPARC heterogeneous refinement, respectively, employing 2-dimer, 3-dimer, 4-dimer, 5-dimer initial models generated from the PADI6 map of CPL. The four classes from CryoSPARC heterogeneous refinement were subjected to non-uniform refinements. Among these, the 3-dimer class and 4-dimer classes yielded reasonable reconstructions and were selected for subsequent 3D classification. To improve the visibility of the third dimer in the 3-dimer assembly, a local 3D classification without alignment was performed with a mask focused on the third dimer. The class exhibiting clear density was selected and subjected to 3D refinement, yielding a 3-dimer reconstruction at an overall resolution of 3.7 Å. The class lacking clear density for the third dimer was refined separately by non-uniform refinement, resulting in a 2-dimer reconstruction with an overall resolution of 3.8 Å.

Similarly, to enhance the density of the third and fourth dimers in the 4-dimer assembly, a local 3D classification without alignment was performed with a mask focused on these regions. The classes displaying clear densities were selected and refined by 3D refinement, generating a 4-dimer reconstruction with an overall resolution of 4.4 Å.

In parallel, the well-resolved 4-dimer class obtained from RELION 3D classification was used to additional rounds of 3D classification. A subsequent local 3D classification without alignment was performed, focusing on the fourth and fifth dimers. The classes displaying clear densities for both dimers were selected for 3D refinement, yielding a 5-dimer reconstruction at an overall resolution of 7.9 Å.

### COLDER colorimetric assay for PADI6 activity

PADI6 enzymatic activity was determined using the COLDER colorimetric method^79^, which detects citrulline formation. The reaction mixture (55 μL) contained 50 nM PADI enzyme, 10 mM CaCl₂, and 10 mM benzoyl-L-arginine ethyl ester (BAEE; Aladdin, B105947) in 50 mM HEPES buffer (pH 7.5) supplemented with 150 mM NaCl. Reactions were incubated at 37 °C for 0–240 min and terminated by adding 5 μL of 500 mM EDTA (final 50 mM).

The color-developing reagent was freshly prepared by mixing Solution A and Solution B (1:3, v/v). Solution A contained 80 mM diacetyl monoxime (DAMO; Macklin, D807426) and 2 mM thiosemicarbazide (TSC; Aladdin, T492440) in deionized water, while Solution B contained 3 M H₃PO₄, 6 M H₂SO₄, and 2 mM (NH₄)₂Fe(SO₄)₂•6H_2_O (Aladdin, A112650). The mixed COLDER reagent was kept at 4 °C in the dark and used within 1 h.

For color development, 200 μL of COLDER reagent was added to each reaction, followed by incubation at 95 °C for 15 min. After cooling to room temperature, absorbance at 540 nm was measured. A standard curve was established using 0–400 μM L-citrulline (Sigma-Aldrich, C7629) standards treated in parallel. PADI6 served as the test group, with PADI1 and bovine serum albumin (BSA; Aladdin, A116563) as positive and negative controls, respectively.

### Resin-embedded mouse oocyte sections for TEM

The ovary tissue from *Ooep^+/^*⁻ and *Ooep⁻^/^⁻* mice were fixed in 2.5% GTH in phosphate-buffered saline (PBS) directly. Oocytes samples after microinjection of siRNAs were washed three times with 0.1% bovine serum albumin (BSA) in phosphate-buffered saline (PBS) and fixed in 2.5% GTH in PBS overnight at 4 °C. After fixation, oocytes were embedded in 2% agrose gels were washed three times with PBS and fixed in 2.5% GTH overnight at 4 °C. After fixation, samples were washed three times with 0.1 M PBS and post-fixed with 2% osmium tetroxide (Electron Microscopy Sciences, 19152) for 2 h at room temperature. The samples were then washed three times with PBS and dehydrated through a graded ethanol series (50%, 70%, 80%, 90%, 95%, and 100%) for 20 min at each step.

Dehydrated samples were infiltrated sequentially with 50%, 75%, and 100% embedding resin, followed by embedding in a resin mixture composed of Embed 812 (Electron Microscopy Sciences, 14900), NMA (19000), DDSA (13710), and DMP-30 (13600). Polymerization was carried out at 65 °C for 48 h.

Ultrathin sections (∼50 nm) were prepared using a Leica EM UC7 ultramicrotome (Leica Microsystems, Vienna, Austria) and stained with 4% uranyl acetate (Electron Microscopy Sciences, 22400) and 2.5% lead citrate (Electron Microscopy Sciences, 17800) prior to imaging by transmission electron microscopy. The sections were then mounted onto copper grids and imaged using a Talos L120C transmission electron microscope (Thermo Fisher Scientific).

### In vitro cRNAs synthesis

The *Flag-Ooep* or *Flag-Padi6* plasmids were linearized with the appropriate restriction endonuclease and used as templates for in vitro transcription with either the SP6 or T7 mMESSAGE mMACHINE Kit (AM1340 or AM1344; Invitrogen, Thermo Fisher Scientific, Waltham, MA, USA). The transcribed cRNAs were polyadenylated with approximately 200–250 bp poly(A) tails using the Poly(A) Tailing Kit (AM1350; Invitrogen), precipitated with lithium chloride, and resuspended in nuclease-free water.

### Microinjection of cRNAs or siRNAs

The siRNA oligos were obtained from Genepharma, and a mixture of three oligos targeting one gene were used as follows: siSkp1-forward, 5’-CCUACGAUAAAGUUGCAGATT-3’, siSkp1-forward, 5’-AGACCAUGCUGGAAGAUUUTT-3’ and siSkp1-forward, 5’-AGCGGACAGAUGAUAUUCCTT-3’; siUbe2d3-1-forward, 5’-GGCGCUGAAACGGAUUAAUTT-3’, siUbe2d3-2-forward, 5’-CCUGCUUUAACUAUUUCUATT-3’ siUbe2d3-2-forward, 5’-GUUGCAUUUACAACAAGAATT-3’. For microinjection, GV-stage oocytes were collected in M2 medium supplemented with 2.5 μM milrinone to maintain meiotic arrest. Microinjections were carried out using an NARISHIGE micromanipulator, with ∼10 pL of siRNAs (40 μM) or synthetic *Flag-Ooep* and *Flag-Padi6*cRNAs (350 ng/μL) injected into the ooplasm. Following injection, GV oocytes were cultured in M2 medium with 2.5 μM milrinone at 37 °C in a 5% CO₂ incubator until further use.

### Follicle collection and culture

Follicles were isolated from 14-day-old (ICR) female mice as described previously^80^. Briefly, ovaries were collected in M2 medium and dissociated enzymatically with addition of 1 mg/ml collagenase IV (Thermo Fisher Scientific, 17104019) in M2 medium for a total of 10 minutes at room temperature. Follicles with a diameter of 100 mm with uniformly surrounding granulosa cells were collected for microinjection. Following microinjection, the follicles were transferred to minidrop covered mineral oil in 6-cm dish and incubated in a 1:1 mixture of α-MEM medium (Glutamax, Gibco, 32571028) and F12 Glutamax supplemented with 5% FBS, 1x Insulin-Transferrin Selenium (ITS) (Thermo Fisher Scientific, 41400045), 100 mIU/mL FSH (Gonal-f), 0.1xPenicillin-G/Streptomycin and 200ng/ml GDF9 recombinant protein at 37 °C/5% CO_2_. Half of the medium was exchanged every 2 days. Follicles were fixed for TEM analysis. Growing oocytes were harvested with mechanistic dissection after 7 days of culture.

### Coimmunoprecipitation (co-IP) and LC-MS/MS

GV oocytes after microinjection with *Flag-Ooep* or *Flag-Padi6* cRNAs for 24 h were collected with IP lysis buffer (Beyotime, P0013) and frozen in a –80°C freezer before use. A total of 1000 untreated, *Flag-Ooep* or *Flag-Padi6* overexpression oocytes in every three replicates were lysed on ice for 30 min. After centrifugation at 15000 rpm for 15 min at 4 °C, the supernatant was collected and incubated with anti-Flag magnetic beads (Sigma-Aldrich, M8823) overnight at 4°C. The beads were collected after three washes with IP lysis buffer using a magnetic frame, and the coimmunoprecipitated proteins were washed three times using water and then subjected to for LC-MS/MS.

Bound proteins were reduced (5 mM DTT, 56 °C, 30 min), alkylated (11 mM iodoacetamide, RT, 15 min, dark), and digested with trypsin (1:50, overnight; then 1:100, 4 h) in 100 mM TEAB. Peptides were desalted on C18 SPE columns. Peptides were separated on a reversed-phase column (15 cm, 100 μm i.d.) using a Vanquish Neo UPLC system with a 12-min gradient of 4–99% solvent B (0.1% FA, 80% ACN) at 600 nL/min, and analyzed by timsTOF HT MS in diaPASEF mode (MS scan 300–1500 m/z; MS/MS 395–1080 m/z, 15 m/z isolation window). DIA data were processed with DIA-NN (v1.8) against the *Mus musculus* SP database (17236 entries, plus decoy). Trypsin/P (max 1 missed cleavage) was used; fixed modifications: Met excision and Cys carbamidomethylation. FDR was adjusted to < 1%.

The differential proteins were considered as significant with fold change >1.5 and P value <0.05. The process of GO annotation was performed using the eggnog-mapper software to extract GO IDs from the identified proteins based on the EggNOG database, and then performing functional classification annotation analysis on the proteins according to cellular components, molecular functions, and biological processes. The volcano map and GO visualization were performed using Weishengxin online tools.

### Immunofluorescence staining

Oocytes were fixed in 3.7% paraformaldehyde (PFA) with 0.2% Triton X-100 prepared in PBS for 30 min, followed by permeabilization with 0.5% Triton X-100 in PBS for 20 min. After blocking with 1% BSA in PBST buffer (PBS with 0.1% Triton X-100), the samples were incubated with primary antibodies for overnight at 4 °C. After three times wahing using PBST buffer, oocytes were then sequentially labeled with Alexa Fluor 594– or 488-conjugated secondary antibodies (Thermo Fisher Scientific) and counterstained with 4′,6-diamidino-2-phenylindole (DAPI) for 30 min. Fluorescence images were acquired using an LSM 800 confocal microscope (Zeiss, Oberkochen, Germany). The primary antibodies used in the experiment are as follows: rabbit anti-OOEP (1:100, Invitrogen, PA5-86033), mouse anti-NLRP5 (1:100, Abmart, M034859S), rabbit anti-TLE6 (1:100, Abmart, PU440858S), rabbit anti-SKP1 (1:100, Proteintech, 10990-2-Ig), rabbit anti-UbcH5C/Ube2d3 (1:100, CST, 4330S) and mouse anti-UHRF1 (1:50, Santa Cruz, sc-373750). The secondary antibodies include: Goat anti-rabbit IgG Alexa Fluor 568 (1:200, Thermo Fisher Scientific, A11036), Goat anti-rabbit IgG Alexa Fluor 488 (1:200, Thermo Fisher Scientific, A11034), Goat anti-mouse IgG Alexa Fluor 568 (1:200, Thermo Fisher Scientific, A11031), Goat anti-mouse IgG Alexa Fluor 488 (1:200, Thermo Fisher Scientific, A11029) and Alexa Fluo Plus 647 (1:200, Thermo Fisher Scientific, A30107).

### Proximity ligation assay (PLA)

The detection of endogenous protein interactions between TLE6 and OOEP used Duolink® In Situ PLA® assay (Sigma-Aldrich, DUO92101). Oocytes were fixed in 3.7% PFA in PBST buffer (PBS with 0.2% Triton X-100) at room temperature for 30 min. The subsequent steps were performed according to the manufacturer’s protocol. The rabbit anti-OOEP (Invitrogen, PA5-86033) and mouse anti-TLE6 (Abmart, M034864S) antibodies were used.

### RNA isolation and real-time RT-PCR

Total RNA was isolated from 50 GV oocytes per sample using an RNeasy Mini Kit (Qiagen, 74104) following the manufacturer’s instructions. Then, cDNA was synthesized by reverse transcription using the SuperScrip II Reverse Transcriptase Kit (Invitrogen, 18064014) according to the manufacturer’s instructions. Real-time RT-PCR was performed using SYBR Green PCR Master Mix (Vazyme) on a Real-Time PCR System (Bio-Rad, CFX96). Relative mRNA levels were calculated by normalizing to endogenous *Gapdh* mRNA (internal control). The sequences of the primers for mouse *Skp1*, *Ube2d3* and *Gapdh* were as follows: *Skp1* (GenBank Accession, NM_011543), forward primer: 5ʹ-ATGCCTACGATAAAGTTGCAGAG-3ʹ and reverse primer: 5ʹ-TCCATTCCCAAATCTTCCAGC-3ʹ; *Ube2d3* (GenBank Accession, NM_025356), forward primer: 5ʹ– AACTTAGTGATTTGGCCCGTG-3ʹ and reverse primer: 5ʹ-GGGCTGTCATTAGGTCCCATAA-3ʹ. The primers for the mouse Gapdh were designed as previously described^81^. The experiment was repeated for three independently obtained samples.

### Western blot analysis

Oocytes (100 GV oocytes in each sample) were lysed directly in 1× Laemmli sample buffer (Bio-Rad). The resulting protein samples were denatured at 98°C for 5 min and subjected to separation by sodium dodecyl sulfate‒polyacrylamide gel electrophoresis, followed by gel transfer to polyvinylidene difluoride (PVDF) membranes (Millipore). Next, the membranes were blocked in 5% skimmed milk for 1 h and then incubated at 4°C overnight with the following primary antibodies: The antibodies used in the experiment are as follows: rabbit anti-OOEP (Beyotime, AF0252), mouse anti-NLRP5 (Abmart, M034859S), rabbit anti-TLE6 (Abmart, PU440858S), mouse anti-UHRF1 (Santa Cruz, sc-373750), mouse anti-SKP1 (Proteintech, 67745-1-Ig), rabbit anti-α/β-tubulin (CST, 2148), mouse anti-GAPDH (Proteintech, 60004-1-Ig). Membranes were then incubated at room temperature with goat anti-rabbit or anti-mouse horseradish peroxidase-conjugated secondary antibodies (111-035-003 or 115-035-003, Jackson ImmunoResearch) for 1 h and washed with TBST buffer three times. Finally, the membranes were visualized using an enhanced exposure solution (Millipore, P90720) and a chemical imaging system (BIO-RAD).

### Quantification and statistical analysis

Gene expression dynamics in Figure 1F and Figure S4 were extracted from the processing data of Proteome, Translaome and Transcriptome as published previously^82^. For the RT-PCR, the relative expression level of control group (siNC) was set to 1 and the fold change was calculated in siSkp1 group. The intensities of immunofluorescence images were analyzed using ImageJ (NIH) and ZEN 3.2 (Zeiss) software. The numbers and lengths of CPL fibers were measured from TEM micrographs using ImageJ (NIH). Enzymatic activity assays were performed with five technical replicates. All statistical analyses were performed using GraphPad Prism 9. Error bars in bar charts and line graphs represent standard deviation of the mean. Data in violin plots are presented as median. Statistical significance was determined using an unpaired, two-tailed Student’s t-test for normally distributed data, or the Mann-Whitney test for non-normally distributed data; for three-group comparisons, one-way analysis of variance (ANOVA) with Tukey’s multiple comparison test was used. Statistical significance is indicated as \**p* < 0.05, \*\**p* < 0.01, \*\*\**p* < 0.001, \*\*\*\**p* < 0.0001 and n.s., *p* > 0.05.

## Figures

Figure panels and videos displaying cryo-EM maps or atomic models were generated using UCSF ChimeraX. Maps coloured by local resolution were generated using RELION5. Multiple-sequence alignments and phylogenetic trees were calculated using Clustal Omega^83^. Electrostatic analyses were performed with *coulombic* command and structure contacts were calculated with *contacts* command with default settings in UCSF ChimeraX. Graphs were plotted in GraphPad Prism (GraphPad Software).

## Data availability

Cryo-EM maps has been deposited in the Electron Microscopy Data Bank with the accession numbers EMD-69647 (an asymmetric unit of CPL composite map), EMD-69566 (CPL overall C2 map), EMD-69606 (an asymmetric unit of CPL consensus map), EMD-69605, EMD-69604, EMD-69603, EMD-69601 and EMD-69599 (CPL local refined maps), EMD-69583 (five-helical bundle composite map), EMD-69573 (five-helical bundle consensus map), EMD-69578, EMD-69579, EMD-69582 and EMD-69581 (five-helical bundle local maps), EMD-69608, EMD-69633, EMD-69635, EMD-69637, EMD-69638 and EMD-69639 (recombinant PADI6 dimers). The corresponding refined atomic model of CPLs has been deposited in the Protein Data Bank under accession number 24MC.

## Acknowledgements

We thank the Cryo-EM Facility of Liangzhu Laboratory and Shuimu BioSciences for the support on data collection and computation, and the Core Facility of Liangzhu Laboratory for technical assistance. We thank Xiaolin Tian and Haiteng Deng in the Center of Protein Analysis Technology, Tsinghua University, for XL-MS analysis. This work was supported by the National Key Research and Development Program of China (2025YFA1309900 to M.G., 2022YFC2702300 to Y.Z.), the National Natural Science Foundation of China (NSFC) (32471245 to M.G., U23A20403 to S.Z.), the Zhejiang Provincial Natural Science Foundation of China (LZ24C050001 and LR26C050002 to M.G., LD26H040001 to Y.Z), the Zhejiang Provincial Leading Innovation and Entrepreneurship Team Introduction and Cultivation Program (2024R01024) to M.G.

## Author contributions

M.G. and S.Z. conceived the project. Y.L. prepared cryo-EM samples, collected data and picked particles with the help of S.Y., Y.L., Y.P. and Q.Z.. Q.L. and M.G. calculated the cryo-EM structures. Y.Z. performed most of the functional work with the help of J.J., S.L., M.L., Y.S., Y.Z., and Y.M.. Q.Z. and Y.L. collected the human and mouse variants data. M.G., S.Z., Y.Z. and X.T. designed the experiments. M.G., Y.Z., Y.L. and Q.L. wrote the manuscript with input from all co-authors.

## Declaration of interests

The authors declare no competing interests.

## Figures

**Figure S1.**
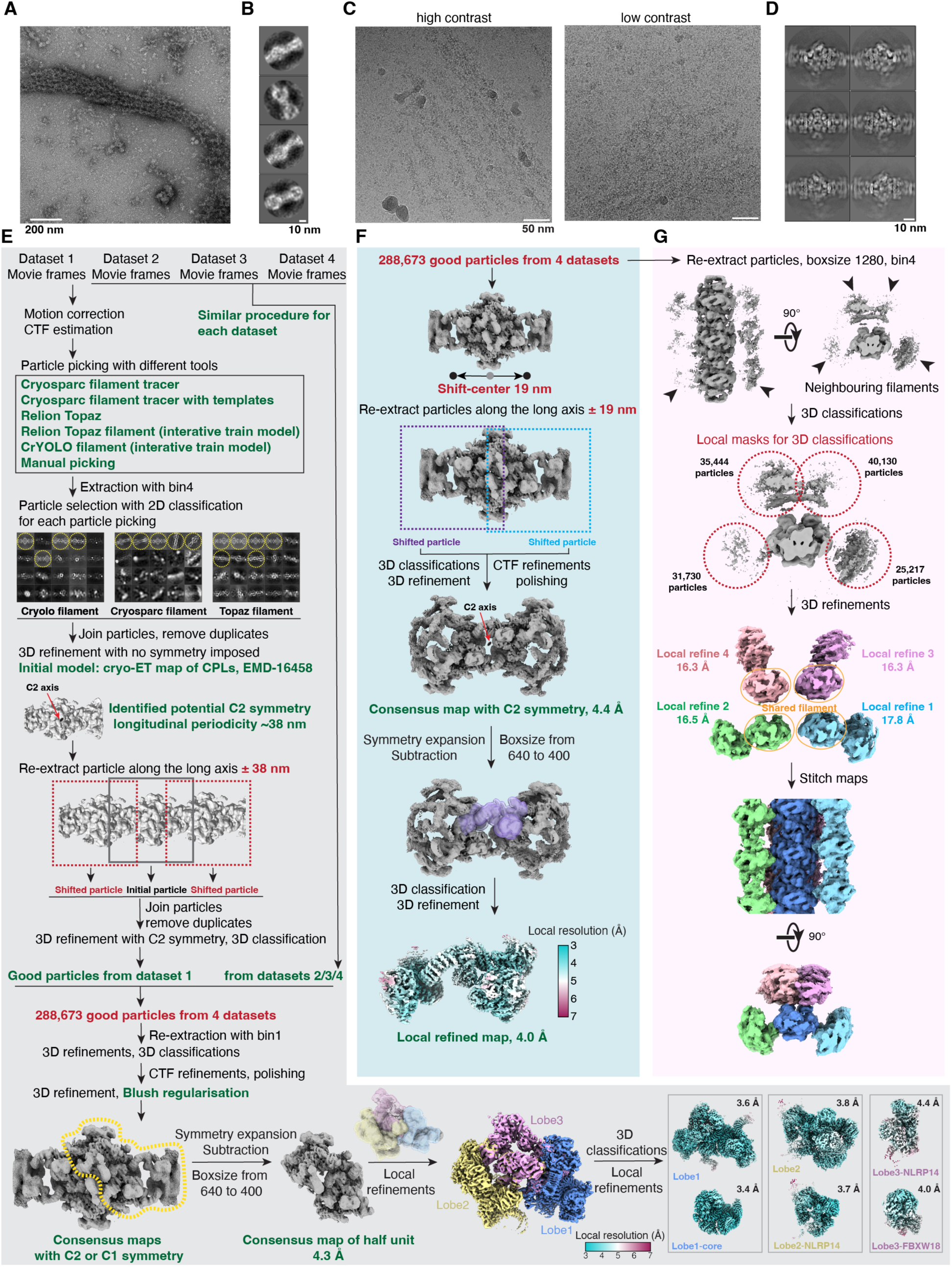
Mouse CPL sample preparation and cryo-EM analysis. (A) Representative negative-stain EM image of CPLs isolated from mouse GV oocytes. (B) Representative 2D class averages of CPLs obtained from negative stain. (C) Representative cryo-EM micrographs of CPLs. (D) Representative 2D class averages of CPLs from cryo-EM data. (E-G) Flow diagram showing the processing strategies for CPLs.

**Figure S2.**
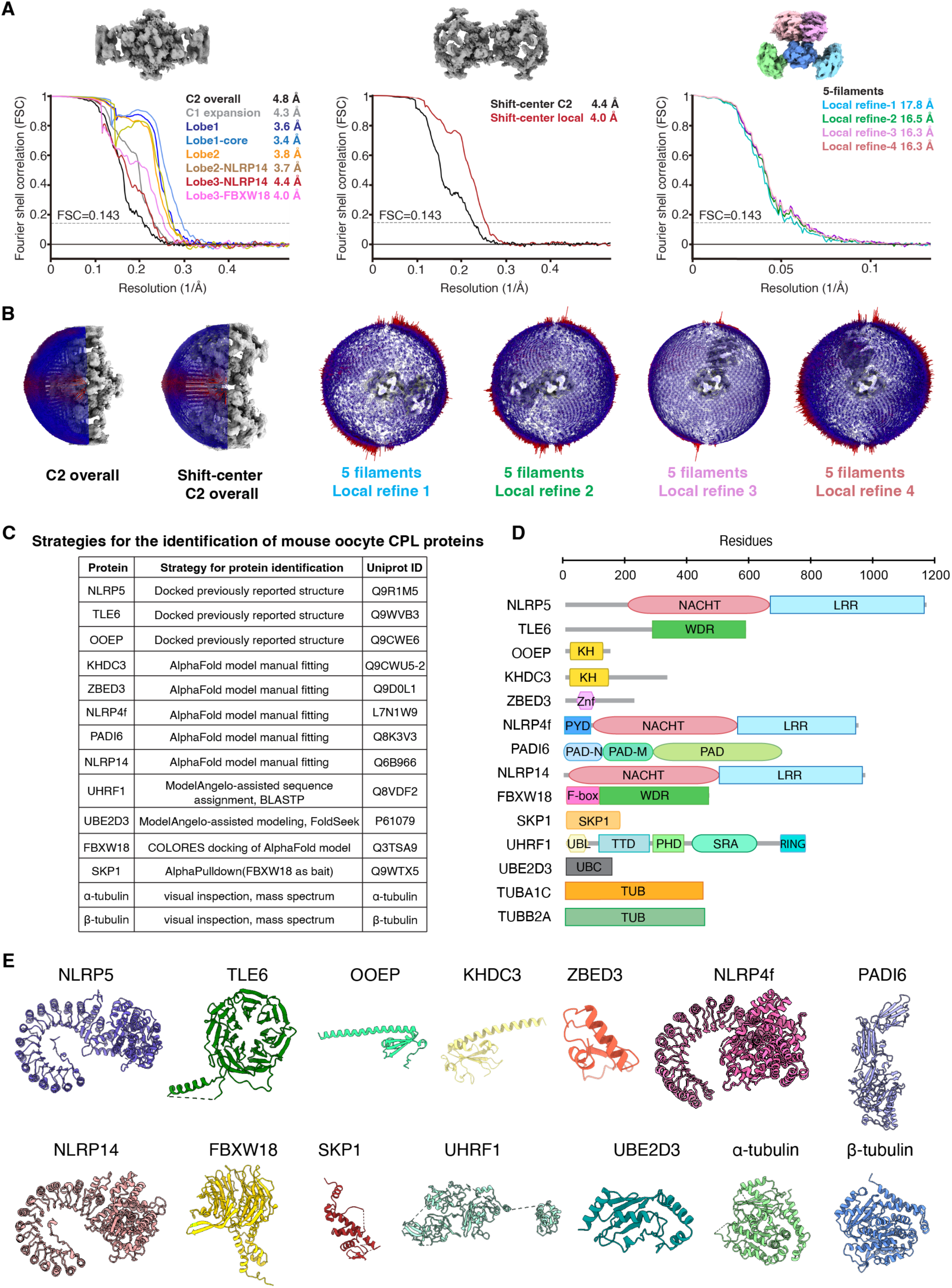
Map resolutions and model building. (A) FSC curves of CPL cryo-EM maps. (B) Particle orientation distributions of CPL cryo-EM maps. (C) Protein assignment strategies for each CPL protein, detailed information are shown in Table S3. (D) Schematic representation of domain architectures of proteins identified in the CPL complex. Protein lengths are shown to scale (residue numbers indicated at top). (E) Atomic models of the CPL proteins, displayed in ribbon representation.

**Figure S3.**
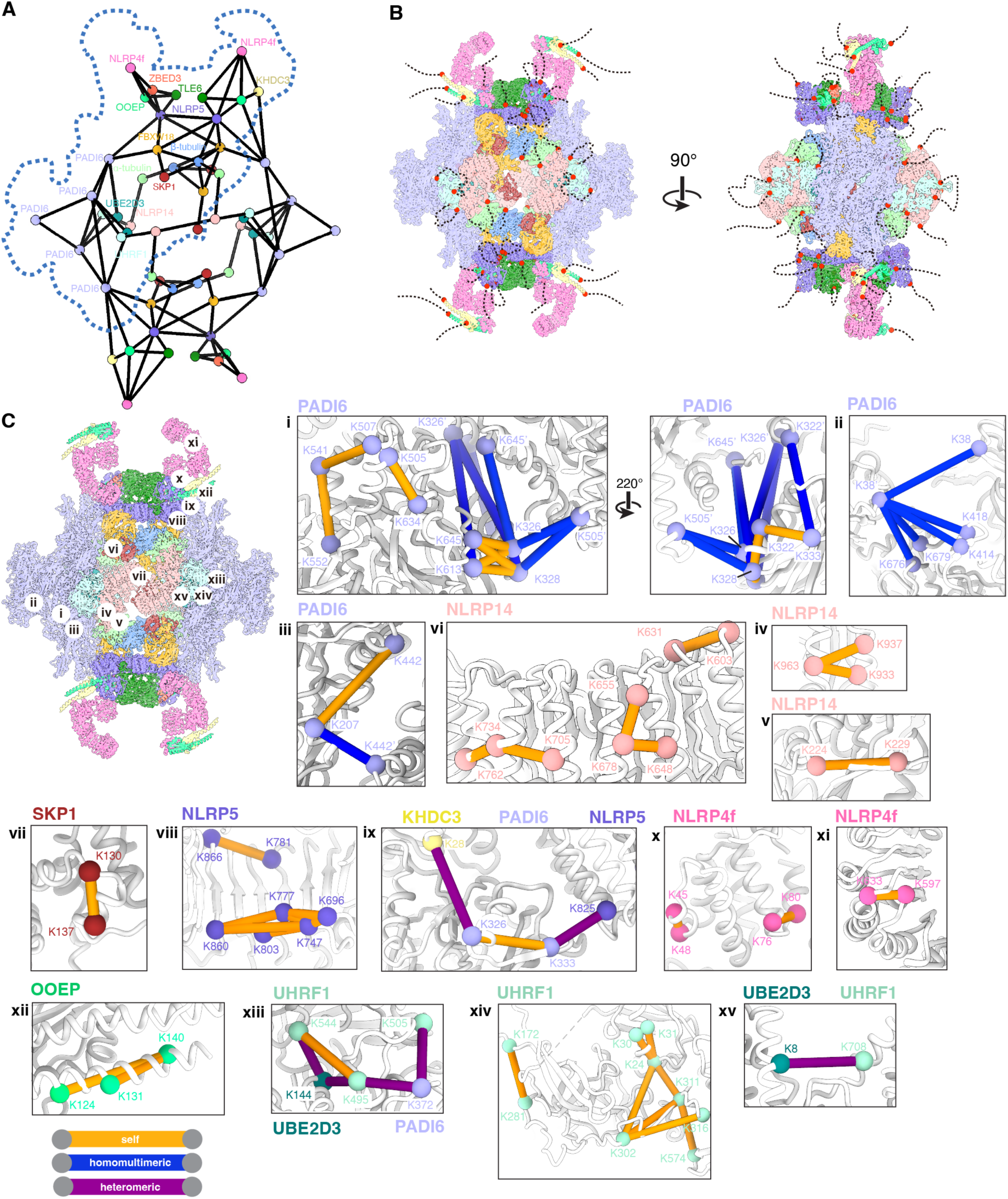
Protein interaction network and XL-MS analysis of mouse CPLs. (A) Protein interaction network within a single CPL repeating unit. (B) The unmodelled loops of the CPL proteins extending outward from the model. (C) The C⍺ atoms of residues cross-linked by BS3 are connected with a line. Orange lines indicate intra-subunit cross-links. Blue lines indicate inter-subunit cross-links between different subunits of the same protein. Purple lines indicate inter-protein cross-links between different proteins.

**Figure S4.**
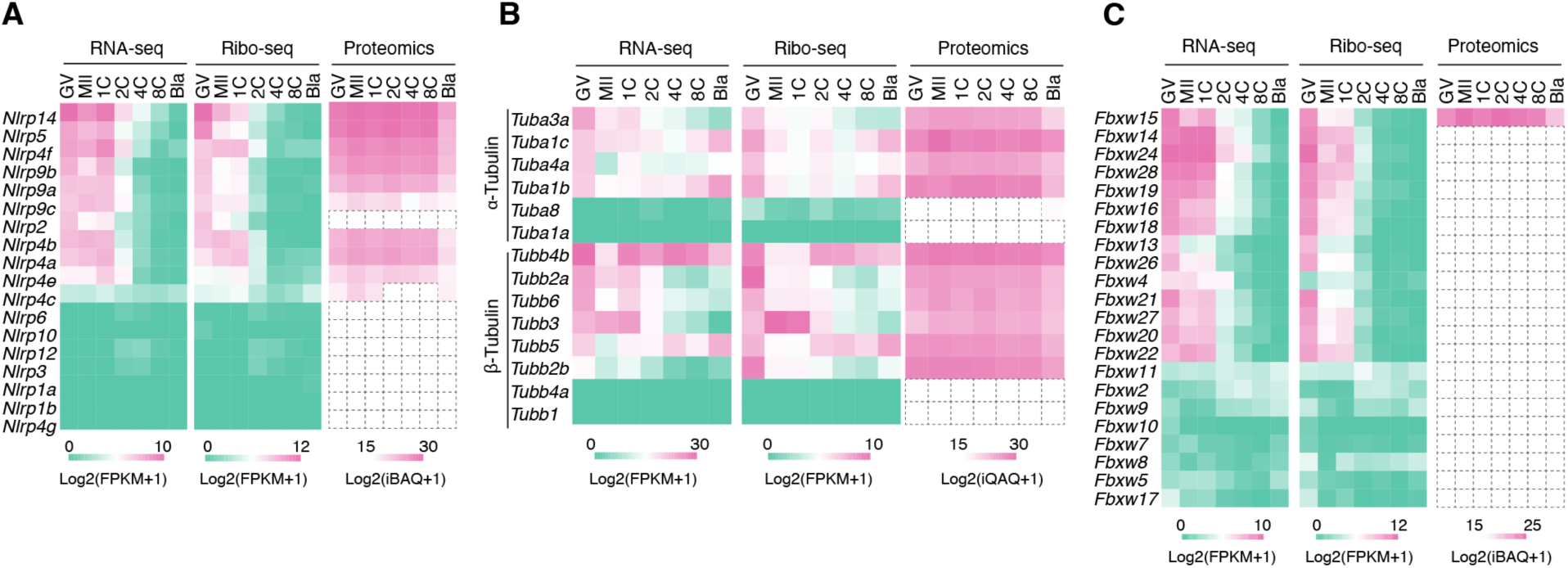
The expression dynamics of NLRP family genes, Tubulins and FBXW family genes. (A) Heat maps showing the dynamics of NLRP family genes based on the mRNA expression level in oocytes and early embryos, with the corresponding translation and protein levels mapped. (B) Heat maps showing the dynamics of 6 genes encoding α-tubulin and 8 genes encoding β-tubulin based on the mRNA expression level in oocytes and early embryos, with the corresponding translation and protein levels mapped. (C) Heat maps showing the dynamics of FBXW family genes based on the mRNA expression level in oocytes and early embryos, with the corresponding translation and protein levels mapped. The dashed box indicates that the proteins were not detected in the mass spectrometry results.

**Figure S5.**
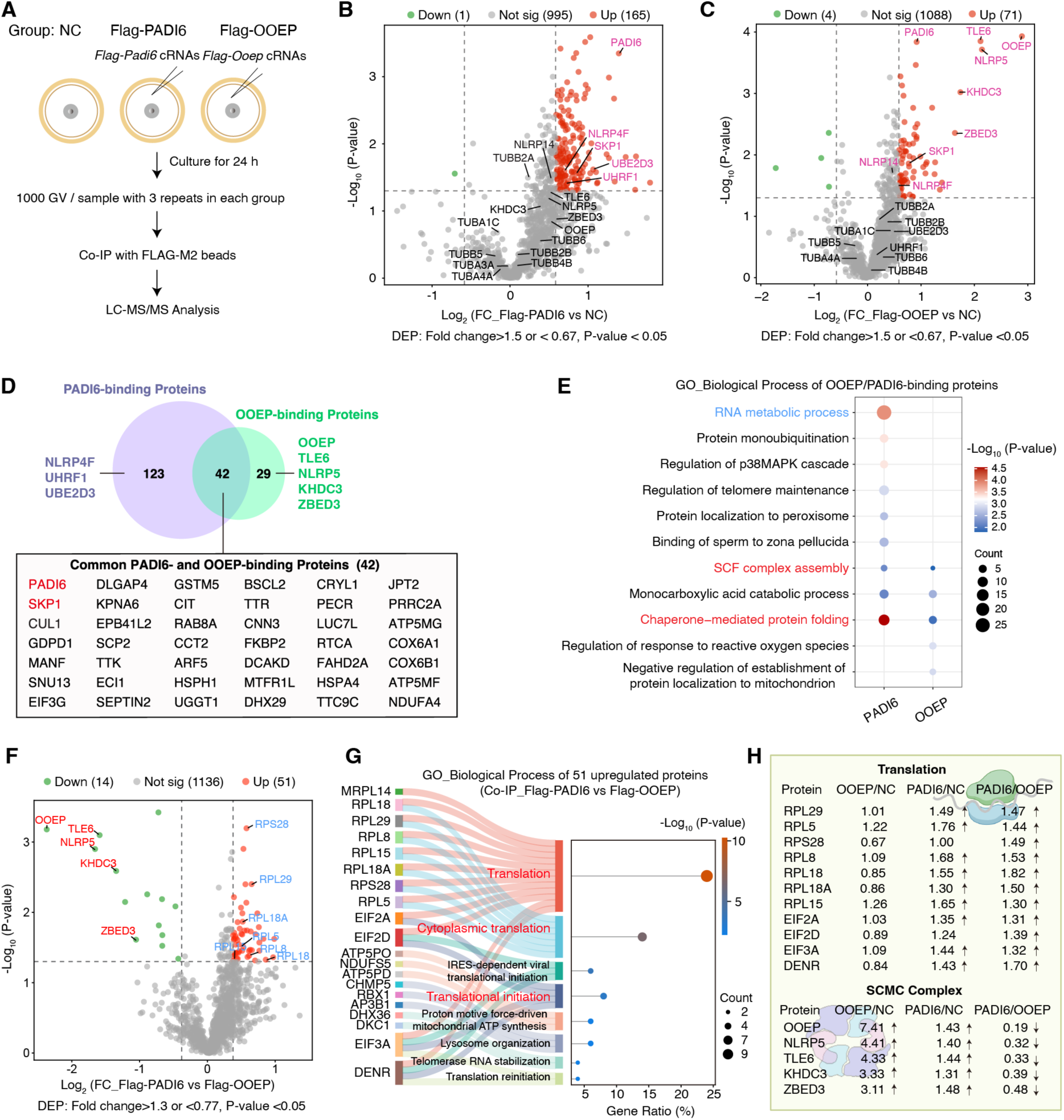
PADI6 and OOEP interact with other CPL components in GV oocytes, revealed by co-IP and LC-MS/MS analysis. (A) Workflow diagram showing the co-IP of exogenous Flag-PADI6 and Flag-OOEP oocyte samples for LC-MS/MS analysis. Three independent biological replicates in each group. (B) Volcano plots showing the significantly enriched proteins (fold change > 1.5, *p*-value < 0.05) in the co-IP of exogenous Flag-PADI6-microinjected GV oocyte sample compared with WT. The CPL components were highlighted in pink with upregulation and marked in black with no change. (C) Volcano plots showing the significantly enriched proteins (fold change > 1.5, *p*-value < 0.05) in the co-IP of exogenous Flag-OOEP-microinjected GV oocyte sample compared with WT. The CPL components were highlighted in pink with upregulation and marked in black with no change. (D) Venn diagram showing the overlap between PADI6– and OOEP-binding proteins. The numbers in each section represent the count of binding proteins that are unique or shared between two co-IP samples. PADI6 and SKP1 are highlighted in red among the overlapping proteins (lower panel). (E) GO analysis of PADI6– and OOEP-binding proteins. (F) Volcano plots showing the significantly differential proteins (DEPs) in the co-IP of microinjection GV oocyte sample of Flag-PADI6 compared with Flag-OOEP group. The SCMC components were highlighted in red with downregulation. The DEPs were defined with Fold change > 1.3 and *p*-value < 0.05. (G) GO analysis showing the enriched biological process of 51 up-regulated proteins in Flag-PADI6 compared with Flag-OOEP. (H) The gene list showing the upregulated proteins in Flag-PADI6 related to translation and downregulated proteins related to SCMC compared with Flag-OOEP.

**Figure S6.**
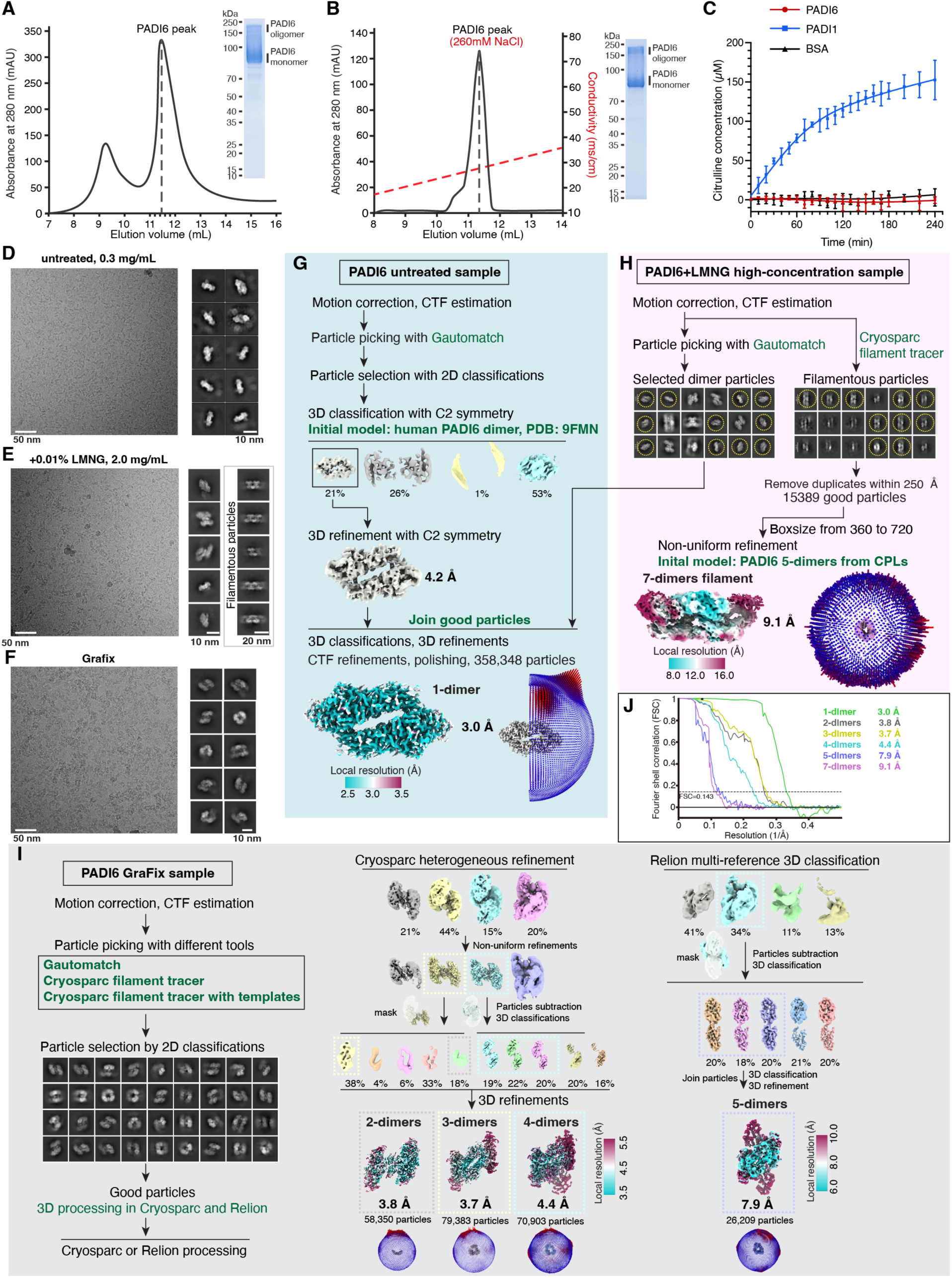
Cryo-EM analysis of recombinant mouse PADI6. (A-B) SEC (Superdex 200 Increase 10/300 GL; A) and ion-exchange (Capto HiRes Q 5/50; B) profiles of mouse PADI6 purified from HEK293F cells, with SDS-PAGE showing a predominant monomeric band and high-molecular-weight bands corresponding to oligomeric PADI6. (C) PADI6 protein-arginine deiminase activity was measured using the COLDER colorimetric assay detecting citrulline formation, with PADI6 (red), PADI1 (blue), and BSA (black) as experimental, positive, and negative controls, respectively. Data are presented as mean ± SD (n = 5). (D) Representative cryo-EM micrographs and 2D class averages of the untreated PADI6 sample. (E) Representative cryo-EM micrographs and 2D class averages of PADI6 in high-concentration LMNG. (F) Representative cryo-EM micrographs and 2D class averages of GraFix PADI6 sample. (G) Flow diagram showing the processing strategy for reconstruction of PADI6 dimer. (H) Flow diagram showing the processing strategy for reconstruction of PADI6 filament. (I) Flow diagram showing the processing strategy for reconstruction of different PADI6 oligomeric states. (J) FSC curves of the PADI6 density maps.

**Figure S7.**
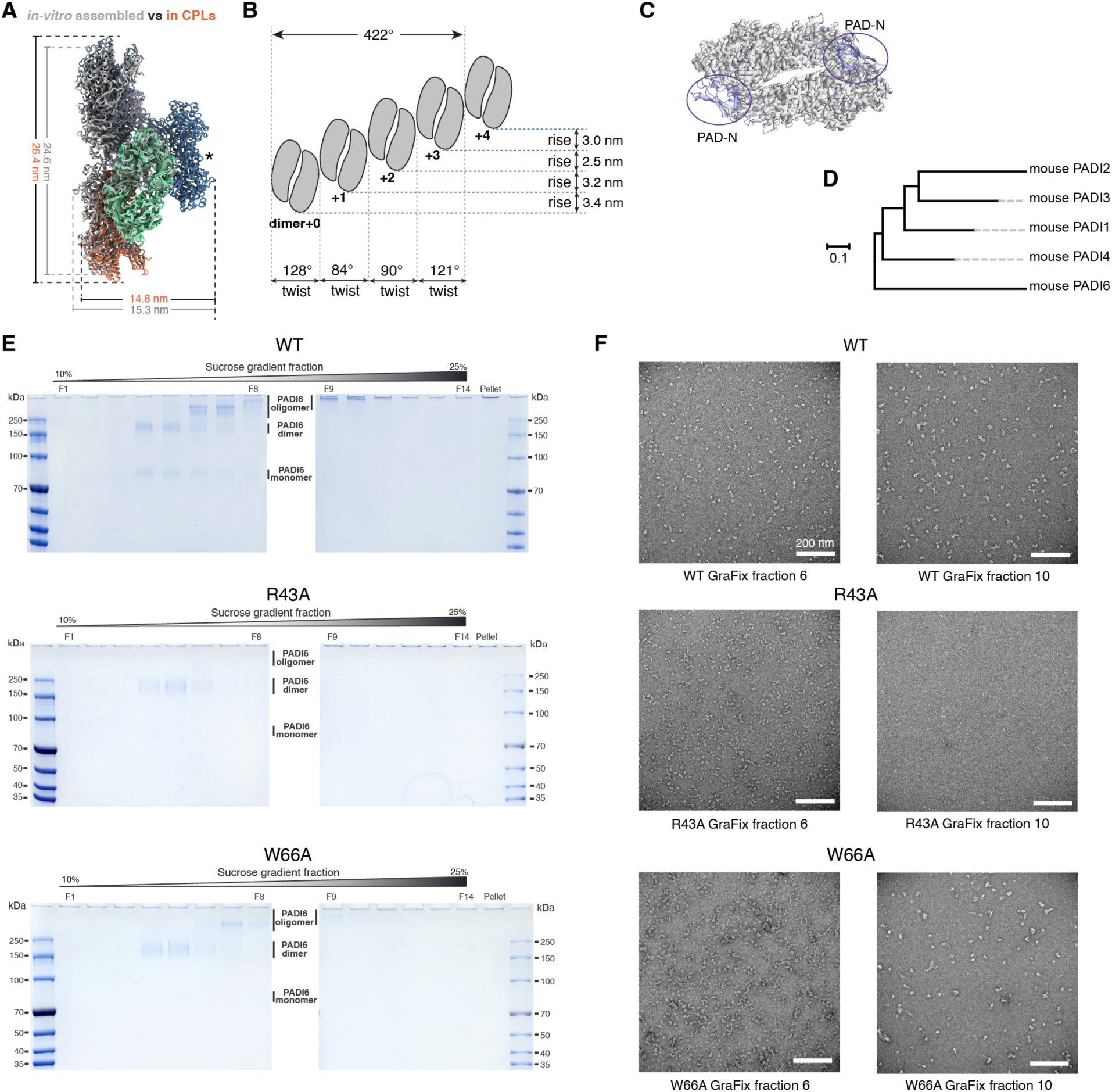
Comparison of GraFix-treated WT PADI6 with R43A and W66A mutants. (A) Structural comparison between PADI6 tubules assembled *in vitro* (grey) and within CPLs. (B) Cartoon schematic illustrating the arrangement of the PADI6 tubules assembled *in vitro*. The helical axis is defined by a line passing through the centroid of the PADI6 tubule and oriented along the longitudinal axis of the CPL. (C) Atomic model fitting of the PADI6 dimer within CPL into cryo-EM map of the *in vitro* assembled PADI6 dimer. PAD-N domain is not visible in the *in vitro* assembled PADI6 dimer. (D) Phylogenetic tree of mouse PADIs. (E) SDS-PAGE of GraFix-treated PADI6 with R43A and W66A mutants across sucrose gradient fractions. (F) Representative negative-stain images of fraction 6 and 10 from GraFix-treated WT, R43A, and W66A PADI6.

**Figure S8.**
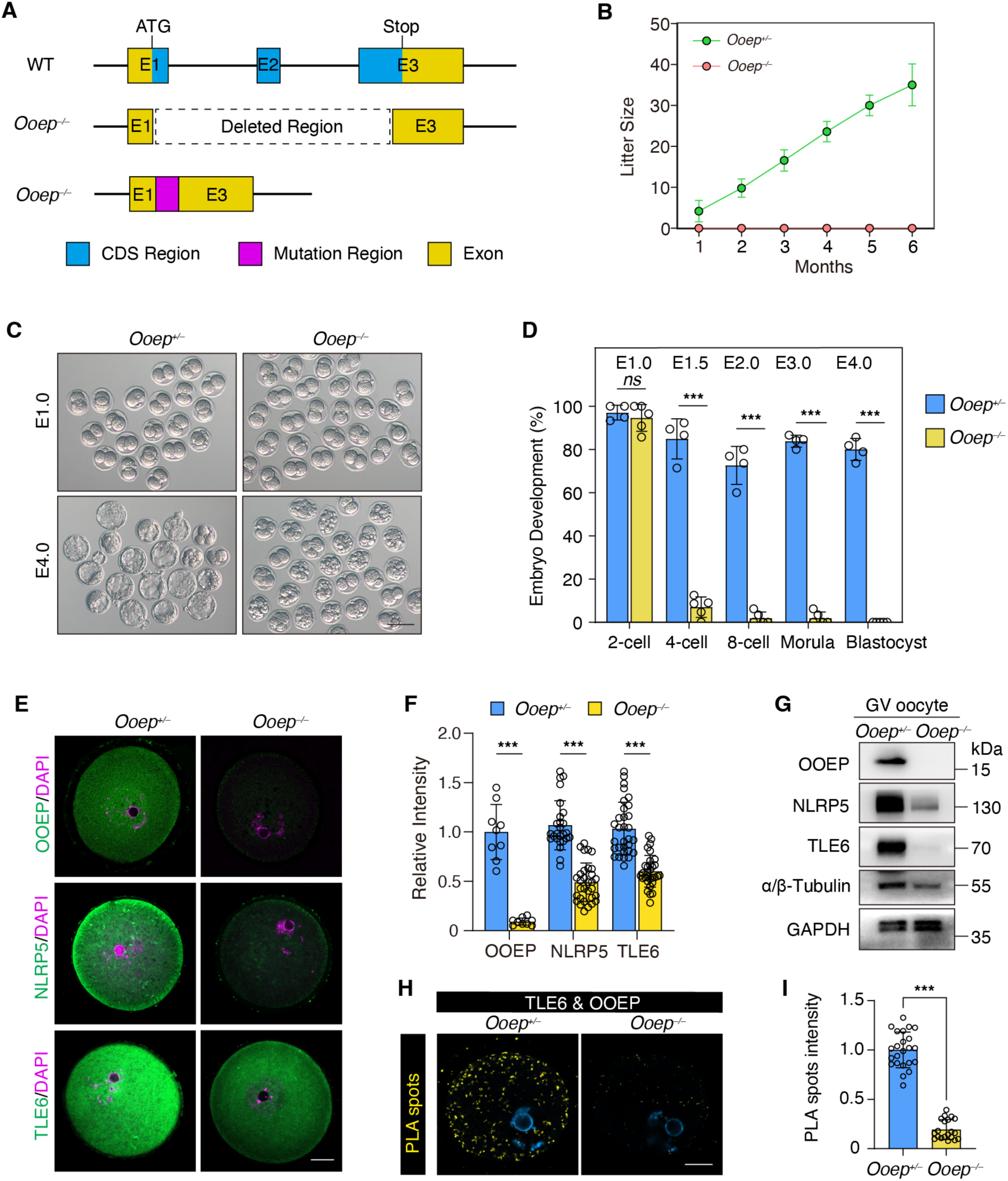
Ooep knockout induces early embryo arrest and SCMC protein instability. (A) Schematic representation of the mouse Ooep gene knockout strategy. (B) Average litter sizes from *Ooep^+/–^* and *Ooep^−/−^* female mice crossed with WT male mice. Data are presented as mean ± SD. (C) Brightfield images of early embryo at E1.0 and E3.0 from *Ooep^+/–^* and *Ooep^−/−^* mice. (D) Quantification of preimplantation embryo development in vitro. The percentages of 2-cell, 4-cell, 8-cell, morula, and blastocyst stages are shown for embryos from *Ooep^+/–^* and *Ooep^−/−^* mice. \*\*\**p* < 0.001; ns, not significant. (E) Immunofluorescence staining of OOEP (green), NLRP5 (green), and TLE6 (green) in GV oocytes from *Ooep^+/–^* and *Ooep^−/−^* mice. DNA is stained with DAPI (magenta). Scale bar: 20 µm. (F) Quantification of immunofluorescence staining intensity of OOEP, NLRP5, and TLE6 in (e). \*\*\**p* < 0.001. (G) Western blot analysis of OOEP, NLRP5, TLE6, and α/β-tubulin protein levels in GV oocyte lysates from *Ooep^+/–^* and *Ooep^−/−^* mice. GAPDH are shown as loading controls. (H) Proximity Ligation Assay (PLA) detecting the interaction between TLE6 and OOEP (yellow spots) in GV oocytes from *Ooep^+/–^* and *Ooep^−/−^* mice. DNA is stained with DAPI (blue). Scale bar: 20 µm. (I) Quantification of PLA spots intensity per oocyte from the results shown in (h). \*\*\**p* < 0.001.

**Figure S9.**
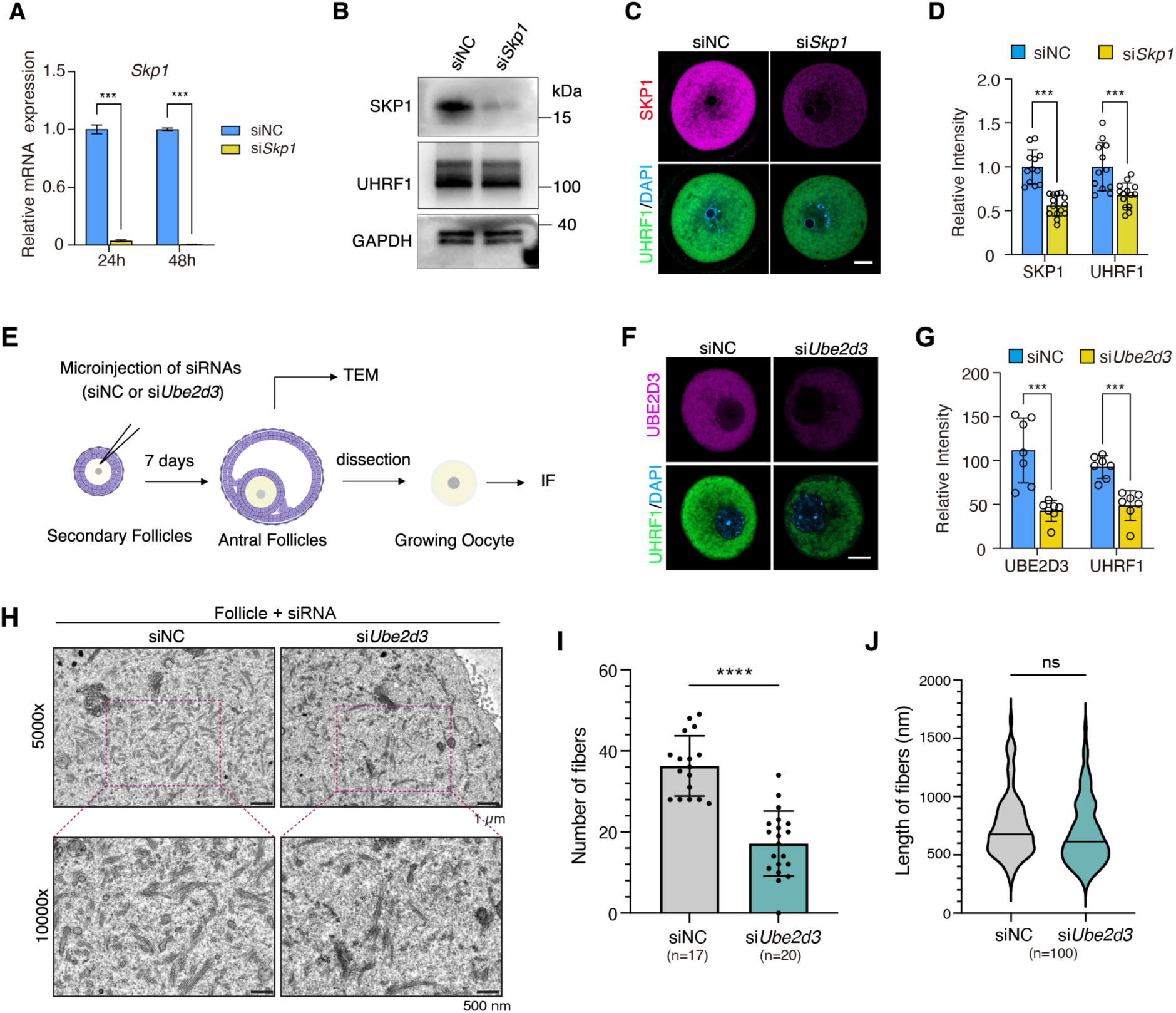
Knockdown of *Skp1* and *Ube2d3* induces UHRF1 protein decrease and CPL filaments loss in oocyte and follicles. (A) Relative mRNA expression of Skp1 in GV oocytes after knockdown with siSkp1 or siNC at 24 and 48 hours. \*\*\**p* < 0.001. (B) Western blot analysis of SKP1, UHRF1, and GAPDH in GV oocytes after knockdown with si*Skp1* or siNC. (C) Images showing immunofluorescence of SKP1 (magenta) and UHRF1 (green) in oocytes from siNC and siSkp1 groups. Scale bar: 20 µm. (D) Quantification of the relative intensity of SKP1 and UHRF1 in oocytes after siSkp1 knockdown. \*\*\**p* < 0.001. (E) The schematic diagram showing the experimental procedure. The siRNAs (siNC or si*Ube2d3*) were microinjected into secondary follicles and antral follicles. After 7 days, oocytes were dissected TEM and immunofluorescence analysis. (F) Representative images showing immunofluorescence of UBE2D3 (magenta) and UHRF1 (green) in growing oocytes from siNC and si*Ube2d3* follicles. Scale bar: 20 µm. (G) Quantification of the relative intensity of UBE2D3 and UHRF1 in growing oocytes after si*Ube2d3* knockdown. \*\*\**p* < 0.001. (H) TEM images showing the morphology of the CPL filaments in follicles from siNC and si*Ube2d3* groups at magnifications of 5000x and 10000x. (I-J) Quantification of the CPL filament number and length in follicles of siNC and si*Ube2d3* groups from 10,000× TEM images (5120 nm × 3840 nm fields of view). \*\*\*\**p* < 0.0001; ns, not significant.

**Figure S10.**
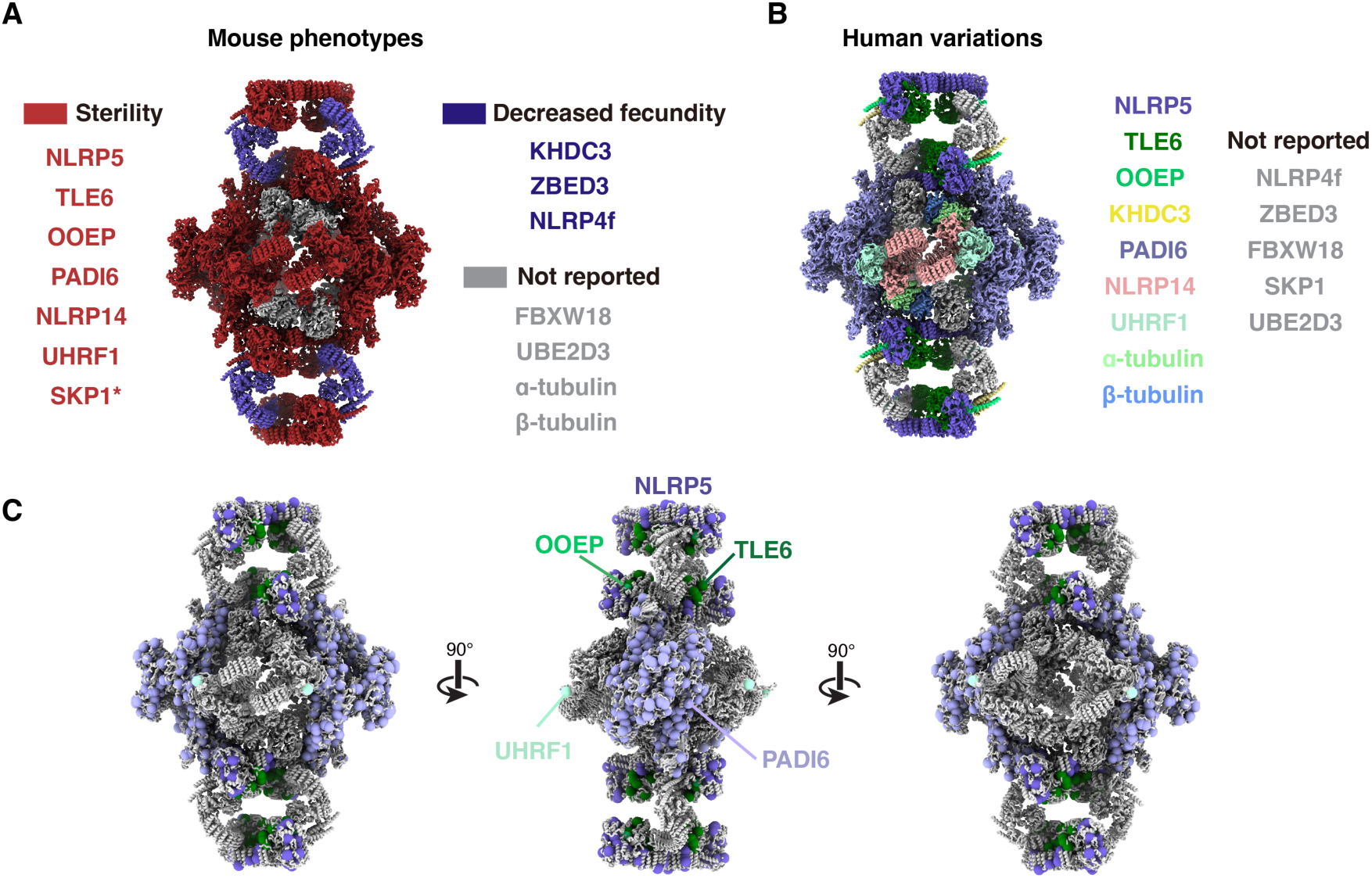
Animal phenotypes and human variants associated with CPL proteins. (A) Proteins associated with mouse female sterility are highlighted in red, whereas proteins associated with decreased fecundity are shown in blue. All other components are rendered in grey. (*) SKP1 KO mice are embryonic lethal and SKP1^cKO^ female mice are sterile. (B) Previously reported human disease-associated CPL proteins are colored coded. All other components are shown in grey. (C) Human missense mutations mapped to the CPL structure, including variants in NLRP5 (31), TLE6 (15), OOEP (1) and PADI6 (31). Variants in *KHDC3L* and *NLRP14* are not shown because no missense mutations were identified. Human variants in *TUBA4A, TUBA1C and TUBB8* have been reported and are not shown in the figure because the tubulin isotypes in CPLs are uncertain.

**Figure S11.**
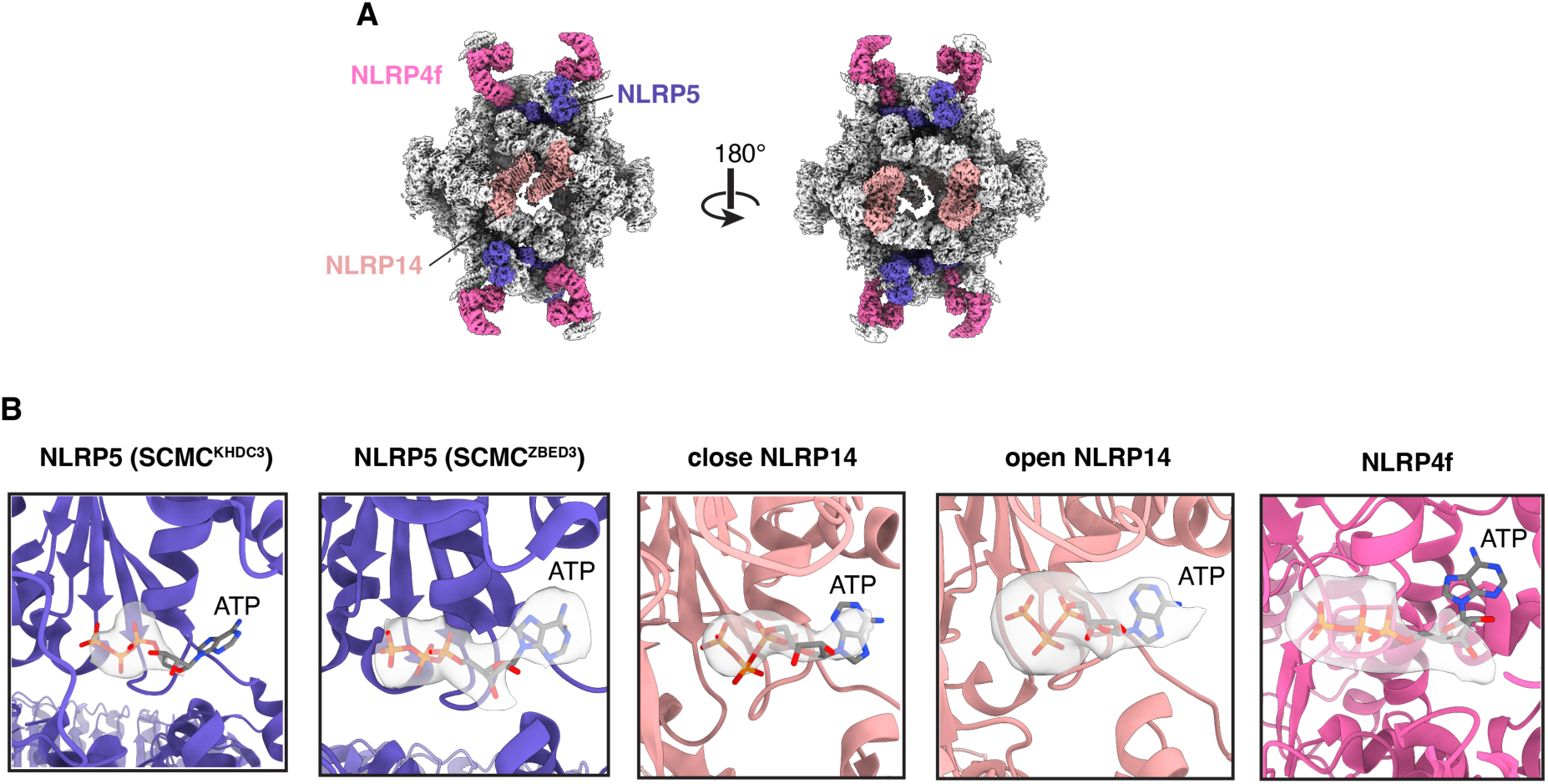
NLRP proteins in mouse CPLs. (A) Locations of NLRP4f, NLRP5 and NLRP14 in CPLs. (B) Cryo-EM densities in the ATP-binding pockets of NLRPs.

## Notes

### Competing Interest Statement

The authors have declared no competing interest.

