## Supplemental Data for "Hierarchical Assembly of Native Cytoplasmic Lattices Revealed by Cryo-EM"

**Tables S1-S7**

**Supplementary Figures Data S1-S11**

**Table S1. Cryo-EM data collection, refinement and validation statistics.**

| | CPL<br>Session 1/3/4 | CPL<br>Session 2 | PADI6<br>1-dimer | PADI<br>2-dimers | PADI6<br>3-dimers | PADI6<br>4-dimers | PADI6<br>5-dimers | PADI6<br>8-dimers filament |
|---|---|---|---|---|---|---|---|---|
| | EMD-69647, PDB 24MC | | EMD-69608 | EMD-69633 | EMD-69635 | EMD-69637 | EMD-69638 | EMD-69639 |
| **Data collection and processing** | | | | | | | | |
| Magnification | 130,000 x | 130,000 x | 130,000 x | 130,000 x | 130,000 x | 130,000 x | 130,000 x | 130,000 x |
| Voltage (kV) | 300 | 300 | 300 | 300 | 300 | 300 | 300 | 300 |
| Electron exposure ($e^{-}$/Å$^{2}$) | 50 | 50 | 50 | 50 | 50 | 50 | 50 | 50 |
| Defocus range (μm) | -1.5 to -2.5 | -1.5 to -2.5 | -1.5 to -2.5 | -1.5 to -2.5 | -1.5 to -2.5 | -1.5 to -2.5 | -1.5 to -2.5 | -1.5 to -2.5 |
| Pixel size (Å) | 0.93 | 0.96 | 0.93 | 0.93 | 0.93 | 0.93 | 0.93 | 0.93 |
| Symmetry imposed | C2 (C1 for local refinements) | | C2 | C1 | C1 | C1 | C1 | C1 |
| Final particle images (no.) | 288,673 | | 358,348 | 58,350 | 79,383 | 70,903 | 26,209 | 15,389 |
| Map resolution (Å) | 3.4 - 4.4 | | 3.0 | 3.8 | 3.7 | 4.4 | 7.9 | 9.1 |
| **Model composition** | | | | | | | | |
| Non-hydrogen atoms | 138,899 | | | | | | | |
| Protein residues | 17,410 | | | | | | | |
| Ligands | 17 (ATP: 5, GTP: 4, $Mg^{2+}$: 4, $Zn^{2+}$: 4) | | | | | | | |
| **Refinement and Validation** | | | | | | | | |
| *B* factors - Protein (Å$^{2}$) | 93.73 | | | | | | | |
| *B* factors - Ligand (Å$^{2}$) | 90.04 | | | | | | | |
| R.m.s.d. - Bond lengths (Å) | 0.007 | | | | | | | |
| R.m.s.d. - Bond angles (°) | 1.184 | | | | | | | |
| MolProbity score | 1.88 | | | | | | | |
| Clashscore | 9.62 | | | | | | | |
| Poor rotamers (%) | 0.74 | | | | | | | |
| Ramachandran plot | | | | | | | | |
| Favored (%) | 94.59 | | | | | | | |
| Allowed (%) | 5.26 | | | | | | | |
| Disallowed (%) | 0.16 | | | | | | | |

**Table S2. Mouse oocyte CPL proteins identified by cryo-EM in this study.**

| Mouse Protein | Alias | Uniprot ID | Human ortholog (Uniprot ID) | Number of residues | Copy per CPL unit |
|---|---|---|---|---|---|
| SCMC | | | | | |
| NLRP5 | MATER, NALP5 | Q9R1M5 | NLRP5 (P59047) | 1163 | 4 |
| TLE6 | | Q9WVB3 | TLE6 (Q9H808) | 581 | 4 |
| OOEP | FLOPED, KHDC2, OEP19, C6orf156 | Q9CWE6 | OOEP (A6NGQ2) | 164 | 4 |
| KHDC3 | FILIA, Filia1.2 | Q9CWU5-2 | KHDC3L (Q587J8) | 346 | 2 |
| ZBED3 | | Q9D0L1 | ZBED3 (Q96IU2) | 228 | 2 |
| NLRP4f | NALP4, PAN2, PYPAF4, RNH2 | L7N1W9 | NLRP4 (Q96MN2) | 937 | 2 |
| Scaffolding proteins | | | | | |
| PADI6 | | Q8K3V4 | PADI6 (Q6TGC4) | 682 | 24* |
| NLRP14 | NALP14, NOD5 | Q6B966 | NLRP14 (Q86W24) | 993 | 4 |
| E3 ubiquitin ligase related | | | | | |
| UHRF1 | ICBP90, NP95, RNF106 | Q8VDF2 | UHRF1 (Q96T88) | 782 | 4 |
| UBE2D3 | UBC5C, UBCH5C | P61079 | UBE2D3 (P61077) | 147 | 4 |
| FBXW18 | | Q3TSA9 | Uncertain | 469 | 5** |
| SKP1 | EMC19, OCP2, SKP1A, TCEB1L | Q9WTX5 | SKP1 (P63208) | 163 | 5 |
| Tubulins | | | | | |
| α-tubulin | | P68373*** | Uncertain | | 4 |
| β-tubulin | | Q7TMM9*** | TUBB8 (Q3ZCM7) | | 4 |

* This includes 2 PADI6 dimers exhibiting flexible densities that are not included in the final atomic model.

** Regions with strong density were confidently assigned to FBXW18. The four remaining segments, although not definitively identifiable, were modeled as FBXW18 to facilitate complete model visualization.

*** Owing to the resolution limitation, the isoforms of α- and β-tubulin could not be resolved. Based on the mass spectrometry results, TUBA1C and TUBB2A, which represent the most likely tubulin species, were used for modeling.

**Table S3. Strategies for the identification of mouse oocyte CPL proteins.**

| Protein | Uniprot ID | Strategy for protein identification |
|---|---|---|
| NLRP5 | Q9R1M5 | 1) Fitted mouse SCMC dimer (NLRP5-TLE6-OOEP, PDB: 8h93) into density map, confirmed by sidechains. |
| TLE6 | Q9WVB3 | 1) Fitted mouse SCMC dimer (NLRP5-TLE6-OOEP, PDB: 8h93) into density map, confirmed by sidechains. |
| OOEP | Q9CWE6 | 1) Fitted mouse SCMC dimer (NLRP5-TLE6-OOEP, PDB: 8h93) into density map, confirmed by sidechains. |
| KHDC3 | Q9CWU5-2 | 1) Manually identified KH domain near OOEP by visual inspection;<br>2) Fitted Alphafold2 model of KHDC3 into the density, confirmed by sidechains.<br>3) Assigned as isoform Filia1.2 because it's more abundant than the canonical isoform Filia1.6 (Ohsugi et al. doi:10.1242/dev.011445). |
| ZBED3 | Q9D0L1 | 1) Found a small density near OOEP and TLE6 in the other SCMC;<br>2) Alphafold3 complex prediction of TLE6-OOEP-ZBED3 fits well in the density, identified ZBED3, confirmed by sidechains. |
| NLRP4f | L7N1W9 | 1) Found a NLRP protein by visual inspection;<br>2) Manually fitted all Alphafold2 predictions of NLRP proteins into the density map;<br>3) Identified NLRP4f based on the N-terminal PYD and landmark sidechains (F4, W12, Y13, L14, R15, F28, W67, Y861, Q892, F910, M911, F933), also confirmed L7N1W9 is the correct NLRP4f isoform in CPLs rather than the canonical sequence (Q66X05). |
| PADI6 | Q8K3V4 | 1) Fitted Alphafold2 prediction of PADI6 dimer into density map;<br>2) Confirmed by sidechains. |
| NLRP14 | Q6B966 | 1) Found a NLRP protein by visual inspection;<br>2) Manually fitted all Alphafold2 predictions of NLRP proteins into the density map;<br>3) Identified NLRP14 based on the lack of N-terminal PYD and landmark sidechains (R31, F39, Y43, F51, Y110, H115, F117, Y118, F132, R211, W215, H343, L362, F363, F381, H385, F389, F390, F394, Y395, H427, L428, M431, R443, R445, R481, F482, H487) |
| UHRF1 | Q8VDF2 | 1) Cropped map submitted to ModelAngelo, obtained a sequence (VIWRWLLSSDSPAPSPNTQEGRDYLEELGLAEKYPDGFLLAEAIKKRKSGKG);<br>2) Submitted to BLASTP, identified UHRF1 as the top hit, confirmed by sidechains. |
| UBE2D3 | P61079 | 1) Cropped map submitted to ModelAngelo, obtained a model with mainchain;<br>2) Submitted to FoldSeek with TM-align mode;<br>3) Identified UBE2D3 as the top hit, TM-score is 0.955, Sequence identity is 40.4%.<br>4) Distinguished with other UBE2 isotypes by landmark sidechains (N7,A14,R125,R131,I137). |
| FBXW18 | Q3TSA9 | 1) Manually identified WD40 Repeat domain by visual inspection;<br>2) Generated a bespoke protein library containing 355 mouse WD40 Repeat-containing proteins by searching "WD40" in Uniprot;<br>3) Run COLORES from the SITUS software to systematically fit the WD40 proteins to the cropped density map;<br>4) Identified FBXW18 as the top hit (correlation coefficient 0.697); |

| | | |
|---|---|---|
| | | 5) Distinguished with other FBXW proteins by landmark sidechains (L5,P6,S7,T58,I60,Y75,T82,C97,T121,T152,T153,N158,A206,A207,I232, R273,N285,K298,Y311,V317,Q329,Y374,D382,I403,S420,R433,Q434,C436,E438,M450,R452,H468). |
| SKP1 | Q9WTX5 | 1) Generated a bespoke protein library of candidate CPL proteins;<br>2) Run AlphaPulldown for FBXW18 and candidate CPL proteins;<br>3) Identified SKP1 as SKP1 and FBXW18 form a stable complex in AlphaPulldown prediction, confirmed by sidechains. |
| α-tubulin | | 1) Manually identified tubulin dimers by visual inspection;<br>2) Confirmed the GTP-$Mg^{2+}$ binding state by density map;<br>3) Tentatively assign as TUBA1C (P68373). |
| β-tubulin | | 1) Manually identified tubulin dimers by visual inspection;<br>2) Confirmed the GTP-$Mg^{2+}$ binding state by density map;<br>3) Tentatively assign as TUBB2A (Q7TMM9). |

**Table S4. Custom protein library for cross-linking mass spectrometry analysis.**

**Table S5. Results of cross-linking mass spectrometry analysis.**

**Table S6. CPL proteins associated with human disease or animal phenotypes.**

**Table S7. Previously reported human disease-associated mutations in identified CPL proteins.**

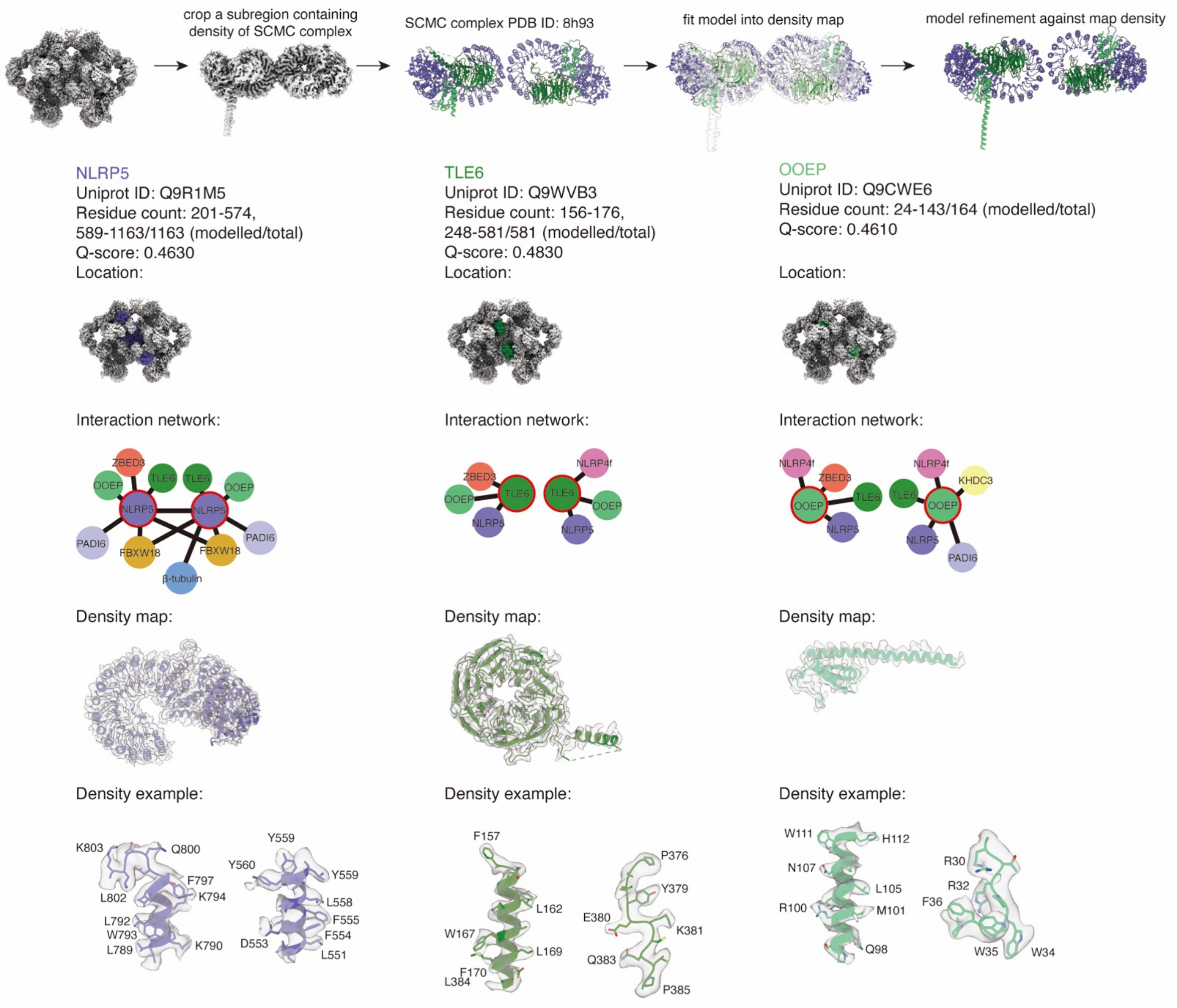


**Data S1. Protein identification and assessment report for NLPR5, TLE6 and OOEP.**

NLRP5, TLE6 and OOEP were assigned based on docking the previously reported structure of the SCMC complex containing these proteins (PDB: 8H93). The assignment was further confirmed by clear sidechain densities.

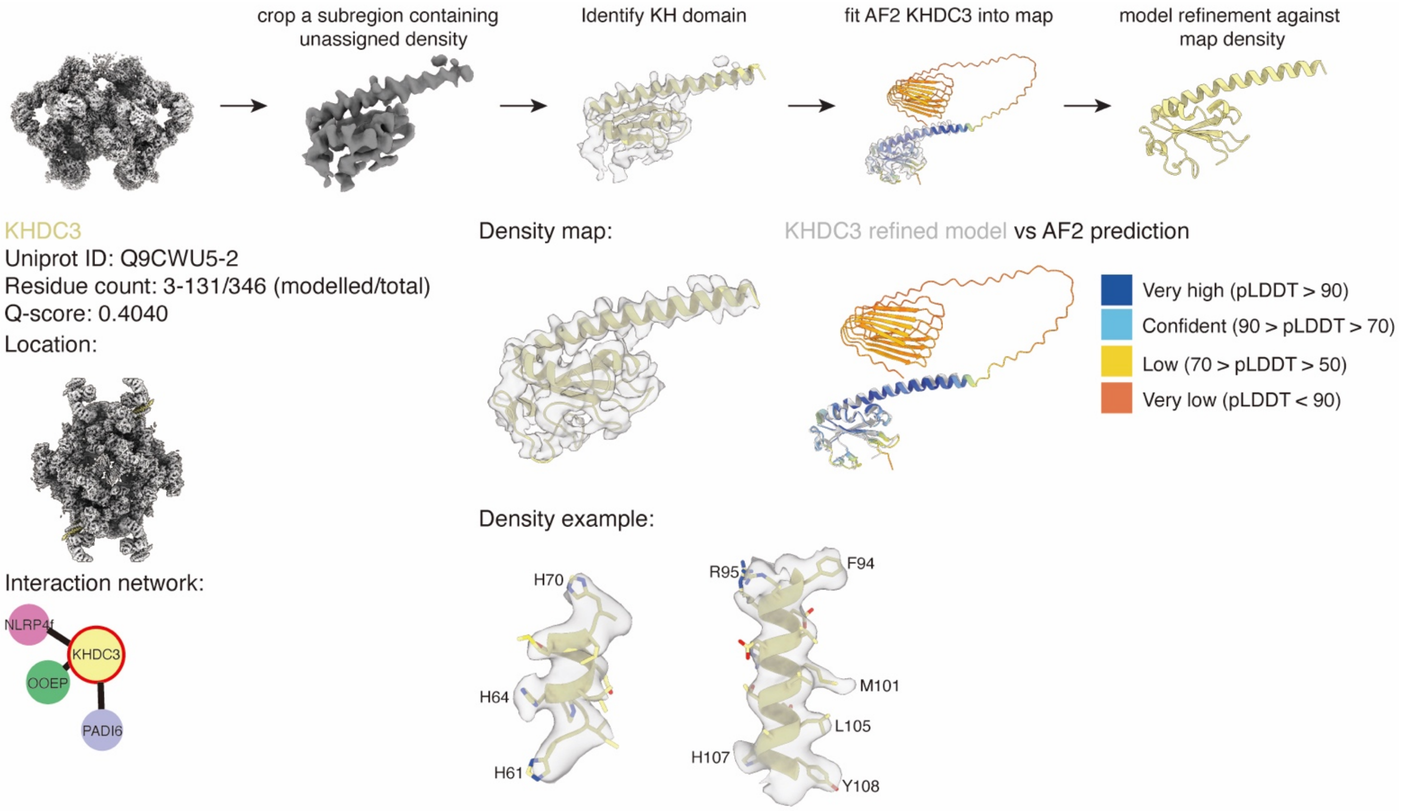


**Data S2. Protein identification and assessment report for KHDC3.**

KHDC3 was assigned by manually identifying the KH domain adjacent to OOEP density, followed by docking the AlphaFold2 model of KHDC3 into the map, which was further validated by well-resolved sidechain densities. The protein was annotated as isoform Filia1.2 due to its higher abundance compared to the canonical isoform Filia1.6.

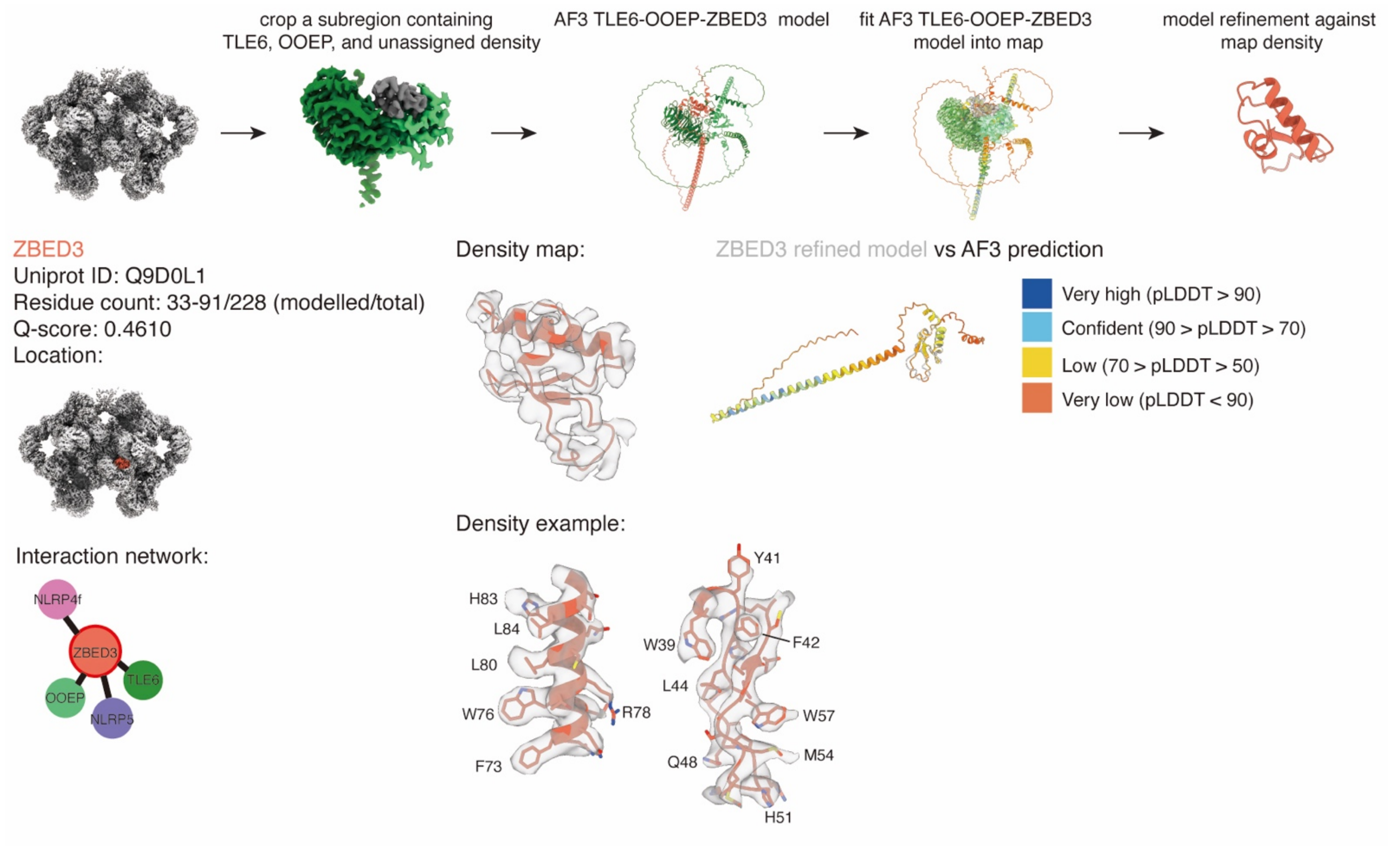


**Data S3. Protein identification and assessment report for ZBED3.**

ZBED3 was assigned by manually docking AlphaFold3 multimer prediction model of TLE6-OOEP-ZBED3. The identification was further validated by well-resolved sidechain densities.

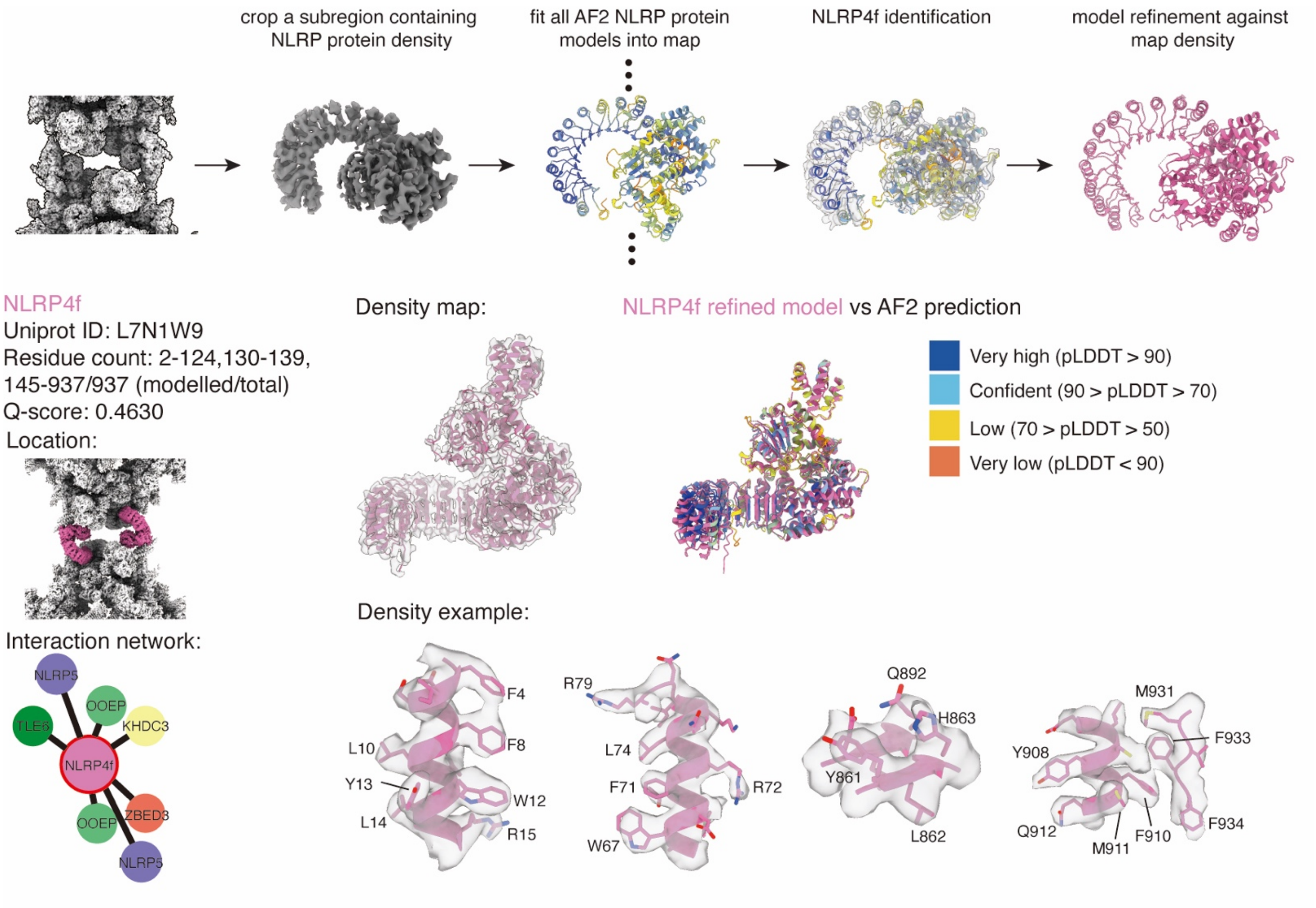


**Data S4. Protein identification and assessment report for NLPR4f.**

NLRP4f was assigned by manually fitting all AlphaFold2 models of NLRP proteins into the unassigned region with clear NLRP protein feature. The NLRP4f was identified by the N-terminal PYD domain and landmark sidechains.

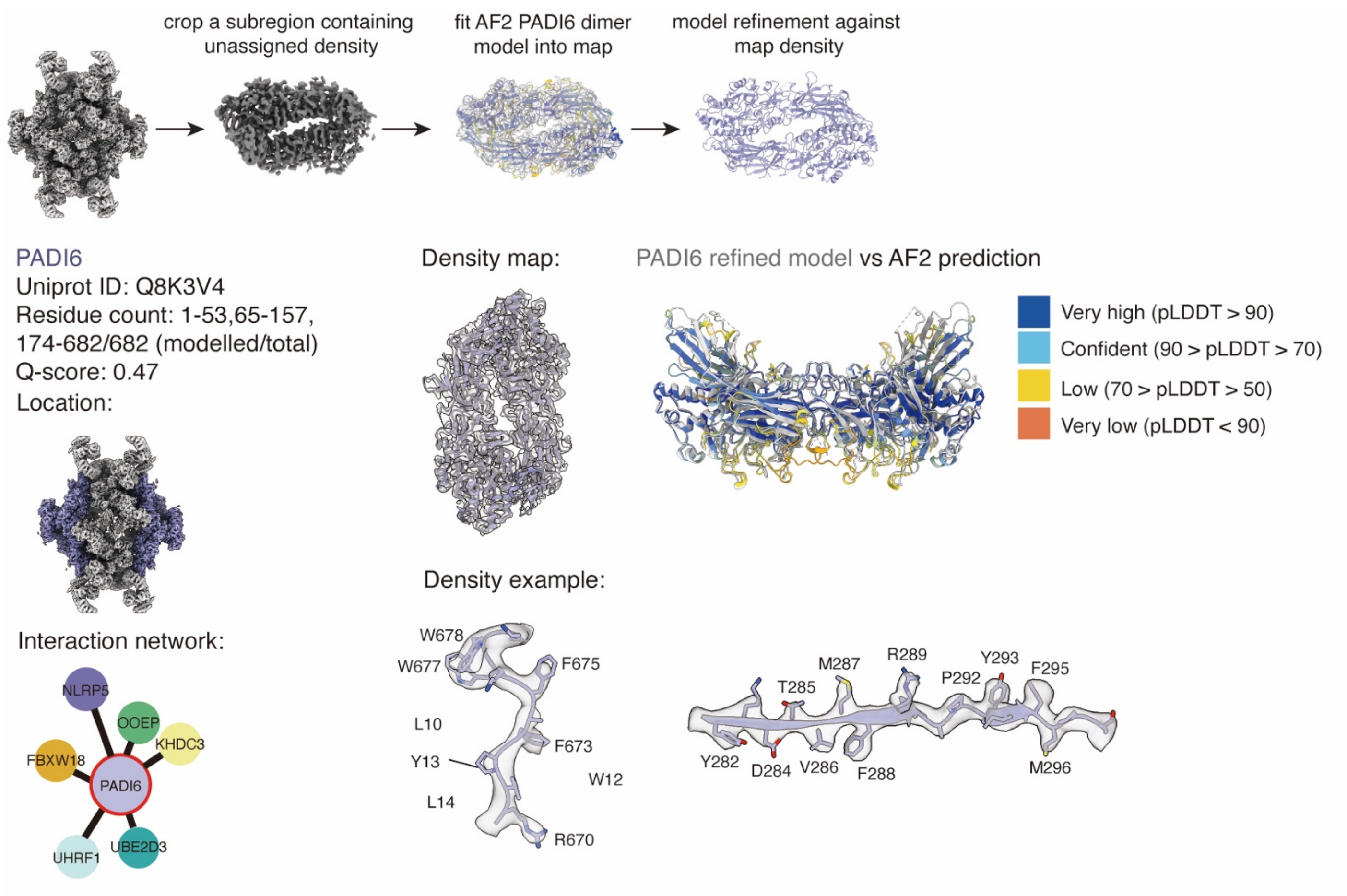


**Data S5. Protein identification and assessment report for PADI6.**
PADI6 was assigned by manually fitting Alphafold2 predicted model of PADI6 dimer into the corresponding densities. The identification was confirmed by clear sidechain densities.

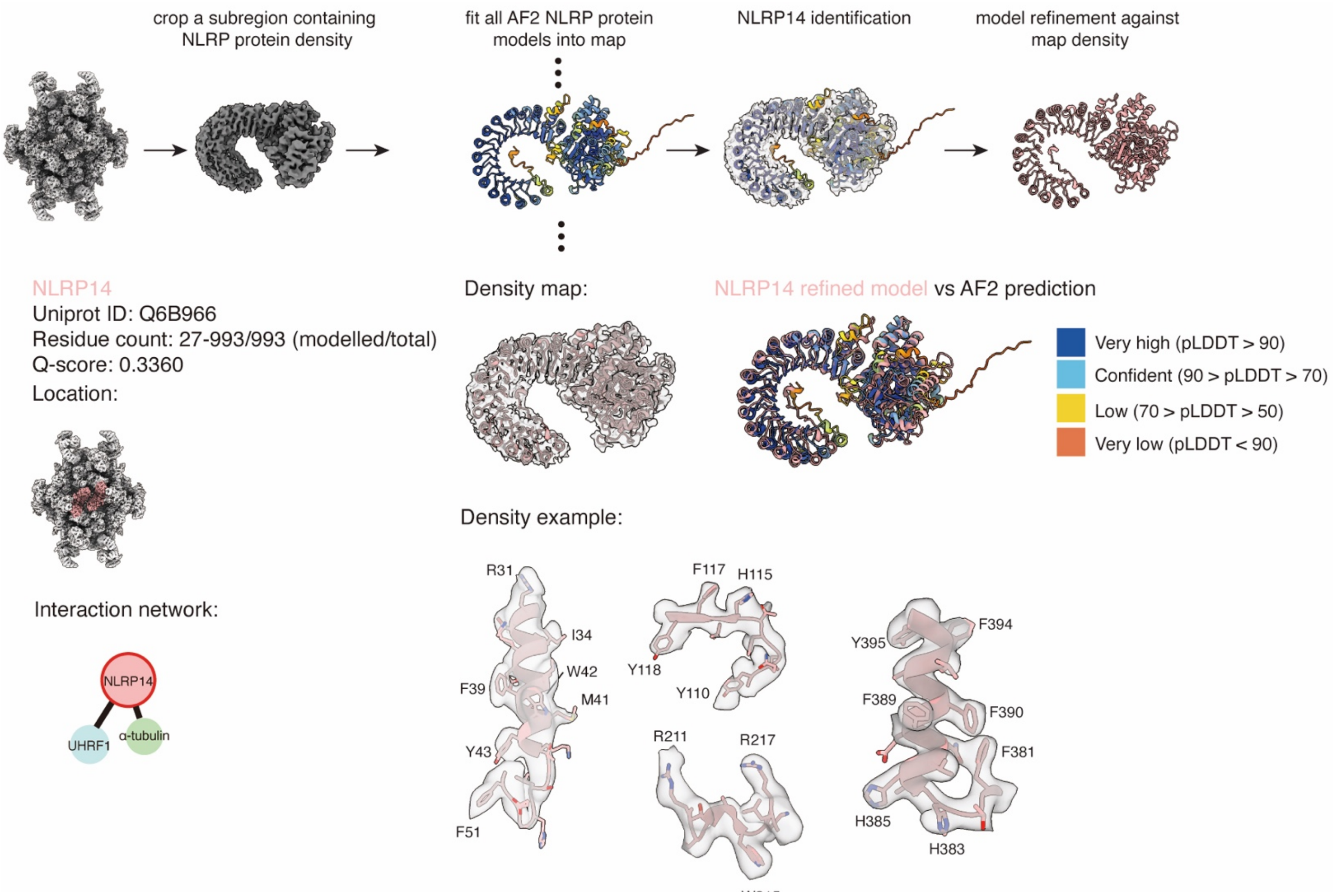


**Data S6. Protein identification and assessment report for NLRP14.**

NLRP14 was assigned by manually fitting all AlphaFold2 models of NLRP proteins into the unassigned region with clear NLRP protein feature. The NLRP14 was identified by the lack of N-terminal PYD domain and landmark sidechains.

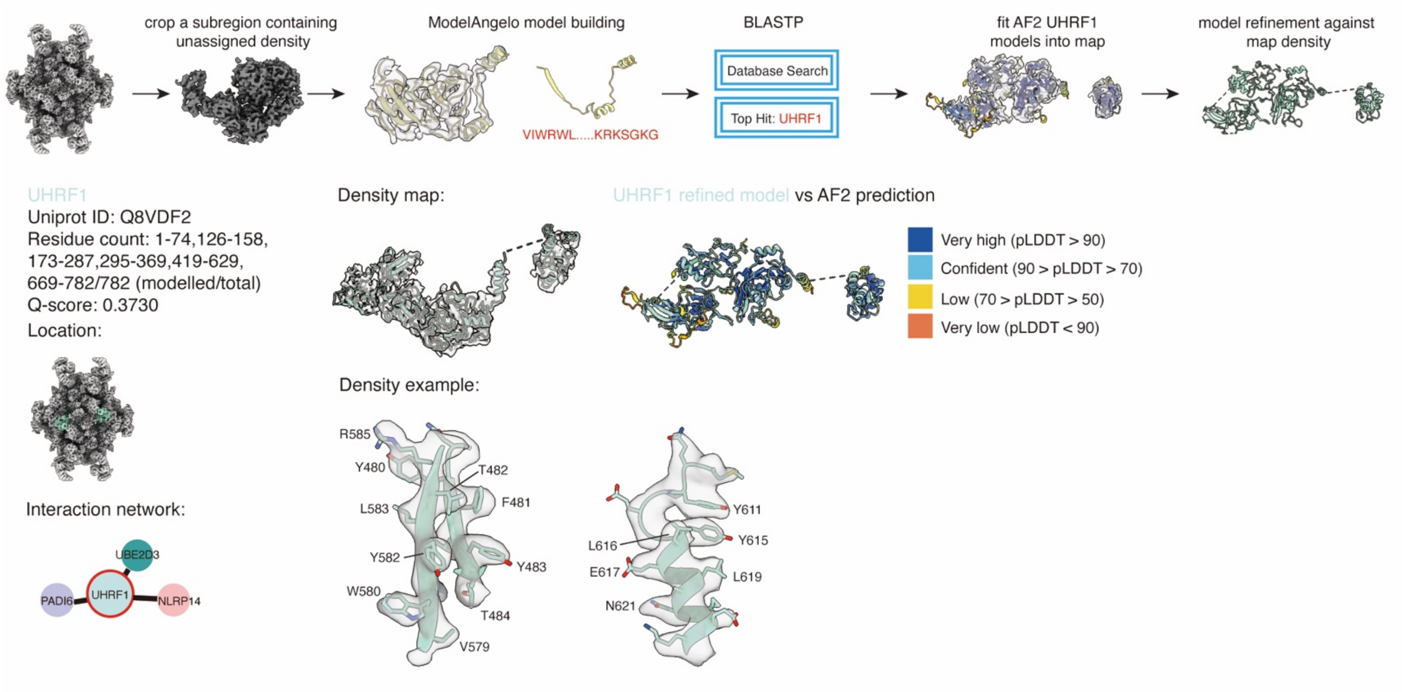


**Data S7. Protein identification and assessment report for UHRF1.**

UHRF1 was assigned based on a specific sequence segment automatically traced by ModelAngelo. The traced sequence was subjected to a BLASTP search against the mouse protein database, and the top hit was identified as UHRF1. Subsequently, the AlphaFold2 model of UHRF1 was fitted into the density map, showing good agreement with the map features. The assignment was further validated by well-resolved sidechain densities.

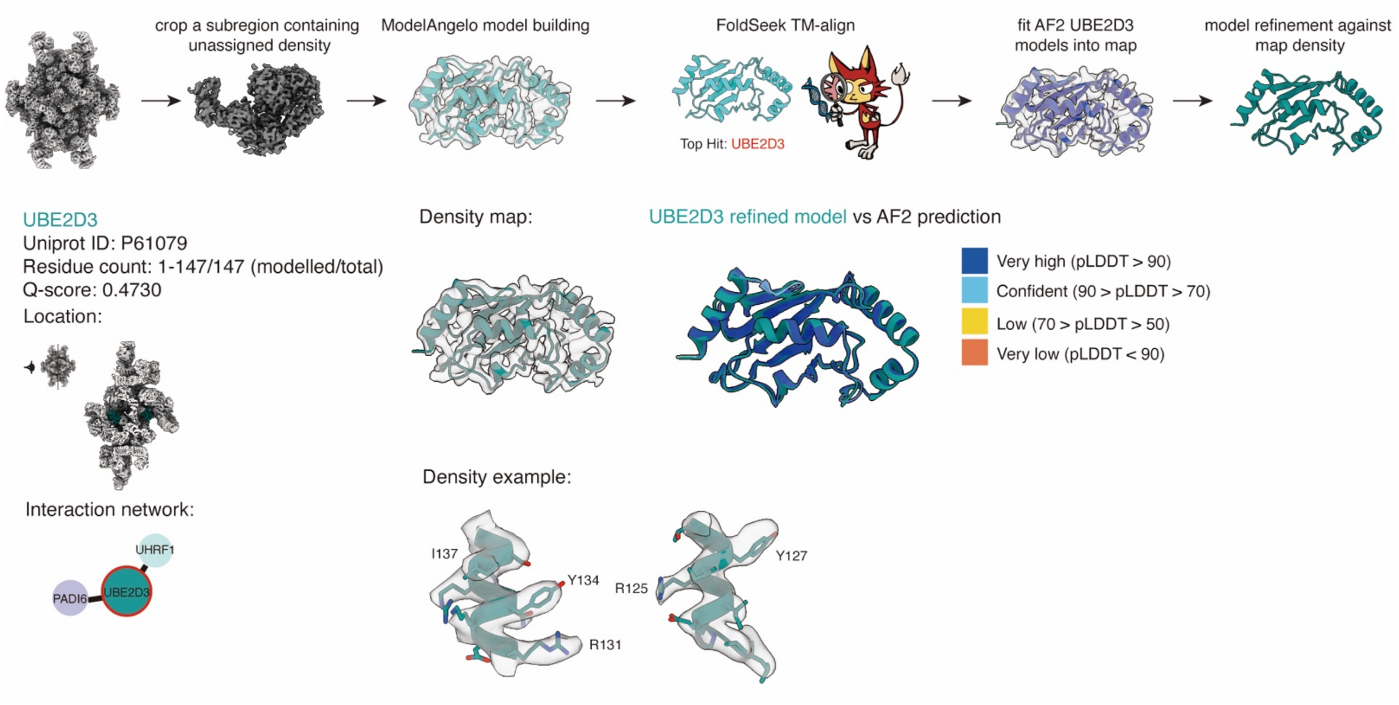


**Data S8. Protein identification and assessment report for UBE2D3.**

UBE2D3 was assigned based on an automated trace by ModelAngelo. The mainchain model was subjected to FoldSeek server using TM-align mode, which identified UBE2D3 as the top hit with a TM-score of 0.955 and a sequence identity of 40.4%. The specific UBE2 isotype was confirmed by sidechain densities.

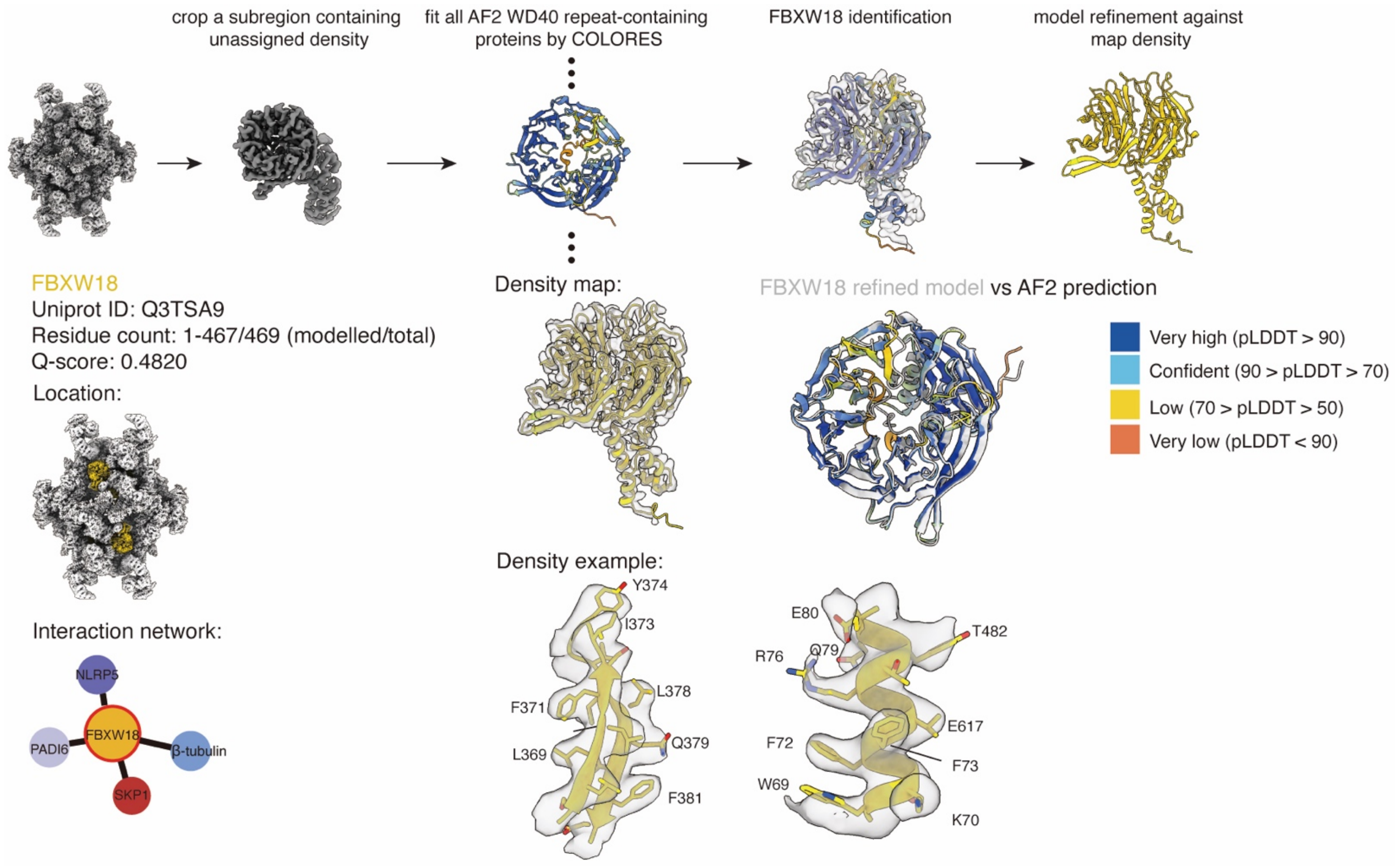


**Data S9. Protein identification and assessment report for FBXW18.**

FBXW18 was assigned by automatically fitting WD40 domain-containing proteins into unassigned, well-resolved density adjacent to open NLRP14 using COLORES, which identified FBXW18 as the top hit with a correlation coefficient of 0.697. The assignment was further identified by landmark sidechains. In contrast, the other two less well-resolved densities within the CPL core region and adjacent to closed NLRP14 were not insufficient to confidently assign FBXW18.

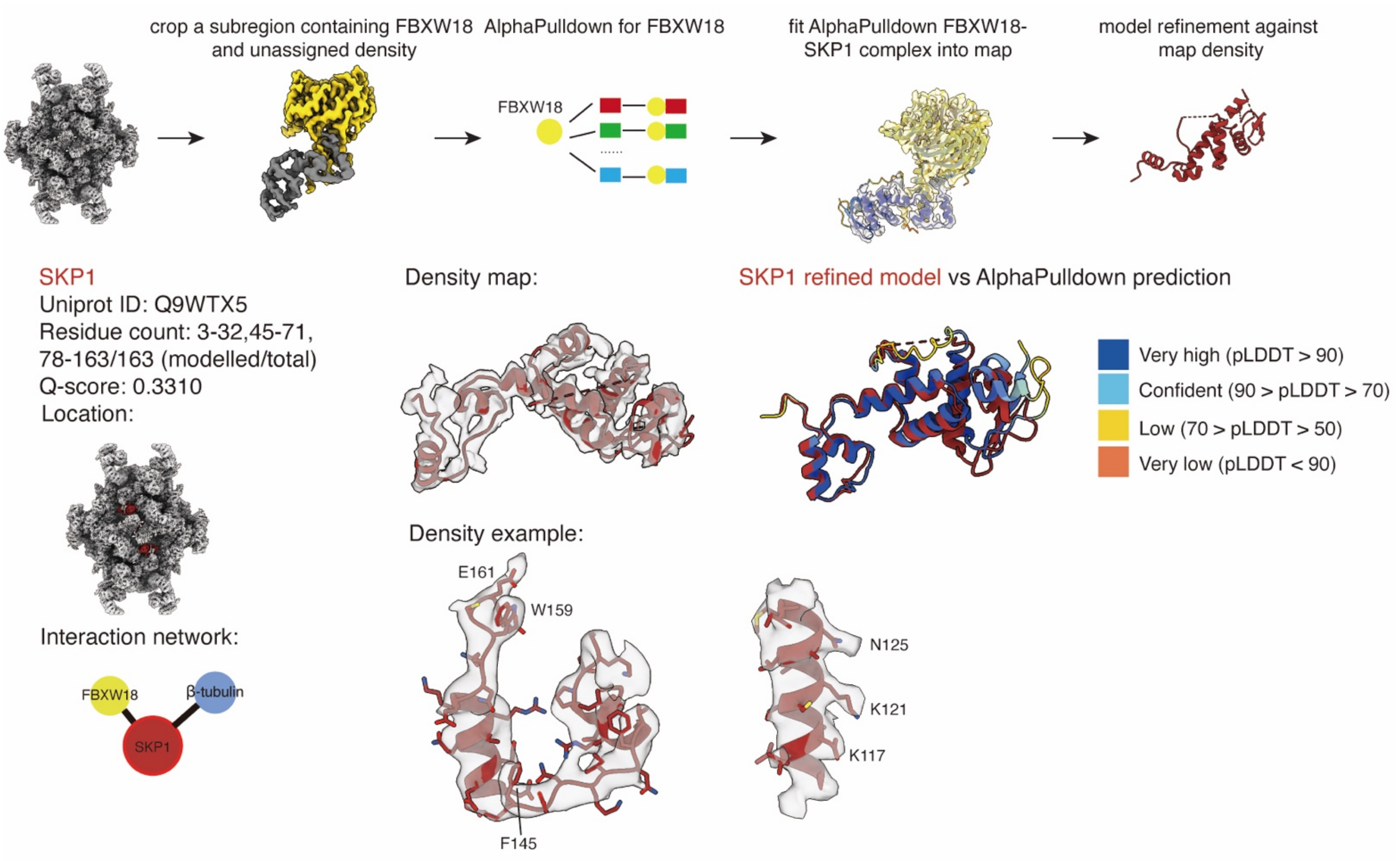


**Data S10. Protein identification and assessment report for SKP1.**

SKP1 was assigned based on AlphaPulldown prediction using FBXW18 as bait, which indicated that SKP1 formed a stable complex with FBXW18. The predicted SKP1-FBXW18 complex was fitted into the corresponding unassigned density and showed good agreement with the map. The assignment was further confirmed by well-resolved sidechain densities.

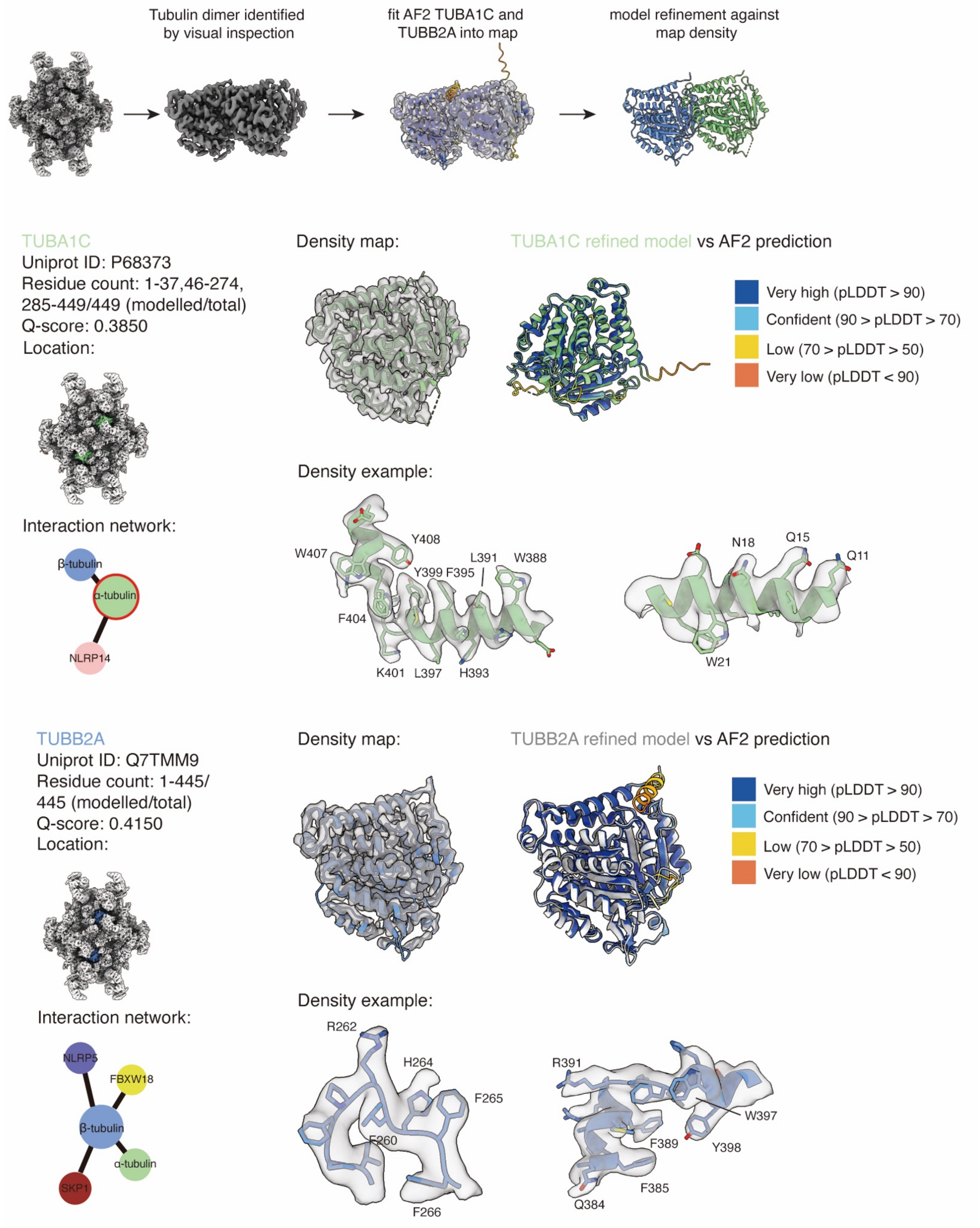


**Data S11. Protein identification and assessment report for TUBA1C and TUBB2A.**
The α- and β- tubulin dimer was assigned by visual inspection of the characteristic tubulin fold and dimeric architecture. Based on the mass spectrometry results, TUBA1C and TUBB2A, which represent the most likely tubulin species, were used for modeling.
